# A B73 x Palomero Toluqueño mapping population reveals local adaptation in Mexican highland maize

**DOI:** 10.1101/2021.09.15.460568

**Authors:** Sergio Perez-Limón, Meng Li, G. Carolina Cintora-Martinez, M Rocio Aguilar-Rangel, M. Nancy Salazar-Vidal, Eric González-Segovia, Karla Blöcher-Juárez, Alejandro Guerrero-Zavala, Benjamin Barrales-Gamez, Jessica Carcaño-Macias, Denise E. Costich, Jorge Nieto-Sotelo, Octavio Martinez de la Vega, June Simpson, Matthew B. Hufford, Jeffrey Ross-Ibarra, Sherry Flint-Garcia, Luis Diaz-Garcia, Rubén Rellán-Álvarez, Ruairidh J. H. Sawers

## Abstract

Generations of farmer selection have produced a unique collection of traditional maize varieties adapted to the environmental challenges of the central Mexican highlands. In addition to agronomic and cultural value, Mexican highland maize represents a good system for the study of local adaptation and acquisition of adaptive phenotypes under cultivation. In this study, we characterized a recombinant inbred line population derived from the cross of the B73 reference line and the Mexican highland maize variety Palomero Toluqueño. Evaluation over multiple years in lowland and highland field sites in Mexico identified genomic regions linked to yield components and putatively adaptive morphological traits. A region on chromosome 7 associated with ear weight showed antagonistic allelic effects in lowland and highland fields, suggesting a trade-off consistent with local adaptation. We identified several alleles of highland origin associated with characteristic highland traits, including reduced tassel branching, increased stem pigmentation and the presence of stem macrohairs. The oligogenic architecture of characteristic morphological traits supports their role in adaptation, suggesting they have arisen from consistent directional selection acting at distinct points across the genome. We discuss these results in the context of the origin of phenotypic novelty during selection, commenting on the role of *de novo* mutation and the acquisition of adaptive variation by gene flow from endemic wild relatives.

## INTRODUCTION

Climatic trends and a need to reduce the level of agronomic inputs have fostered interest in the development of crop varieties that are not only high yielding but *resilient* - *i.e.* their performance is stable in the face of diverse, and potentially unpredictable, environmental challenges. One approach to enhance the cultivated genepool for greater resilience is to explore diversity at the extremes of a crop’s distribution (Emon *et al.*, 2015; Dwivedi *et al.*, 2016; Corrado & Rao, 2017; Sousaraei *et al.*, 2021). Thousands of years of effort and care by the world’s traditional farming communities have generated a rich diversity of landrace varieties, collectively adapted to a far broader ecological range than modern breeding material (Bellon *et al.*, 2018). Crop landraces serve to illustrate the mechanisms whereby plants can adapt to environmental stress as well as representing a valuable source of adaptive variation in their own right.

Strong directional selection imposed by prevailing conditions tends to produce highly specialized forms that perform well in their home environment, but relatively poorer in other locations, a process referred to as *local adaptation*. Local adaptation is defined formally as superior performance of local genotypes in their native environment versus non-local genotypes (Anderson *et al.*, 2013; Mitchell-Olds *et al.*, 2007; Hall *et al.*, 2010; Anderson *et al.*, 2011). Concomitantly, the average performance of a locally adapted variety over a range of environments may be poorer than that of a generalist that maintains a reasonable level of performance in all environments - *stability* in the context of plant breeding. Experimentally, the best demonstration of local adaptation is the *reciprocal transplant* experiment, in which varieties of interest are evaluated in a series of common gardens covering the range of their home environments. Both local adaptation and stability can be described with reference to *genotype x environment interaction* (GEI), *i.e.* the degree to which the relative performance of a given variety compared with others depends on environmental conditions (Juenger, 2013; Assmann, 2013; Scheiner, 1993; El-Soda *et al.*, 2014).

By definition, all varieties will likely suffer reduced performance when challenged by environmental stress. GEI describes variety-specific deviations from the environmental main effect: some varieties suffer more than average, while others are better able to mitigate the impact of the stress. In extreme cases, the relative performance of varieties changes between environments, a scenario referred to as *rank changing* GEI. While stress is often considered with respect to a single sub-optimal factor, the same framework applies equally to the complex pattern of challenges presented by different localities. It can be seen that rank changing GEI underpins local adaptation, as defined above. With the advent of comparative genomics and greater understanding of the physiology and cell biology of environmental responses, it has become feasible to begin to characterize the genetic basis of local adaptation (Lovell *et al.*, 2021). Two principal modes of gene action have been proposed to drive rank changing GEI, namely *conditional neutrality* and *antagonistic pleiotropy*. Under conditional neutrality, a given genetic variant is linked to phenotypic change in some environments but not others. A complementary suite of conditionally neutral loci would, theoretically, be sufficient to generate rank changing GEI. Under antagonistic pleiotropy, the sign of the effect of a given variant changes between environments, *e.g.* a beneficial allele in one environment becomes deleterious in another, with a behaviour at a single variant that directly mirrors the whole genotype pattern of GEI. In practice, both behaviors will typically contribute to GEI and, indeed, classification of any given variant will be specific to the environments under consideration (Fournier-Level *et al.*, 2011). In addition, distinguishing conditional neutrality from antagonistic pleiotropy may be limited by statistical power in any given design. To date, studies of local adaptation in wild barley *Hordeum spontaneum* (Verhoeven et al., 2004; Verhoeven et al., 2008), the annual grass *Avena barbata* (Gardner & Latta, 2006; Latta et al., 2007; Latta et al., 2010), the model plant *Arabidopsis thaliana* (Weinig et al., 2003; Fournier-Level et al., 2011) and monkey flower *Mimulus guttatus* (Hall et al., 2010; Lowry et al., 2009) have predominantly found cases of conditional neutrality. That said, examples of antagonistic pleiotropy do exist, although mostly limited to plant model organisms such as *Arabidopsis thaliana* (Scarcelli et al., 2007; Todesco et al., 2010), monkey flower *Mimulus guttatus* (Hall et al., 2010) and *Boechera stricta* (Anderson et al., 2013). An important consequence of the genetic architecture of local adaptation is the degree to which the specialist is constrained by trade-offs that impose an unavoidable cost of poor performance outside of the home environment. In terms of plant breeding, there are analogous implications with regard to how extensively a given variety can be used and how robust yields will be in the face of unpredictable or changing environmental conditions.

In addition to their intrinsic value, crop landraces provide an excellent system to study local adaptation, especially with regard to the rapid change required over the relatively short time frame of domestication. Landraces are dynamic populations, each with a unique identity shaped by biotic and abiotic stresses, crop management, seed handling and consumer preferences. As such, landraces are the product of both direct and indirect farmer selection, natural selection in the face of the local environment and exchange through traditional seed flow networks (Louette *et al.*, 1997; Cleveland & Soleri, 2007; Mercer *et al.*, 2008; Mercer & Perales, 2019). Typically, they are cultivated under low-input conditions and produce a modest but stable yield (Zeven, 1998; Breseghello & Coelho, 2013; Dwivedi *et al.*, 2016). The sustained association of a given landrace population with a given locality results in local adaptation, in the same way it is seen in wild populations, demonstrable by reciprocal transplantation (Janzen *et al.*, 2021).

Maize (*Zea mays* ssp. *mays*) was domesticated from balsas teosinte (*Zea may*s subsp. *parviglumis*) (Matsuoka *et al.*, 2002), about 9000 years ago, in the basin of the Balsas River in Mexico (Piperno *et al.*, 2007). After domestication, maize dispersed and was successfully established in different environments throughout the Americas and, eventually, across the world. In Mexico alone, 59 different native landraces of maize have been described, grown from sea level to 3400 m.a.s.l., in a range of environments, from semi-desert to regions with high humidity and temperature (CONABIO, 2018). One of the environments colonized by early maize was the central highlands of Mexico. The central Mexican highlands are characterized by low atmospheric pressure and temperature, high UV-B radiation, seasonal precipitation, presence of early frosts and low phosphorus availability due to the volcanic origin of the soil (Bellon *et al.*, 2005; Körner, 2007; Mercer *et al.*, 2008; Espinosa-Calderón *et al.*, 2011; Galván-Tejada *et al.*, 2014). Previous work has highlighted the impact of low temperature on unadapted maize varieties, which when grown at low temperatures and exposed to high light intensity, suffer metabolic lesions in chlorophyll synthesis, leading to increased photodamage and chlorophyll turnover (McWilliam & Naylor, 1967). Interestingly, these cold stress-induced symptoms were not observed in highland maize. Highland maize varieties have to mature and complete the grain filling before the first frosts (Alvarado-Beltrán *et al.*, 2019). In warmer lowland conditions, highland material is precocious, flowering in as little as 40 to 50 days.

In the Mexican highlands, farmers have adapted their management practices to improve their chances of obtaining a successful harvest (CIMMYT & Bjarnason, 1994; Eagles & Lothrop, 1994). To maximize the length of the growing season, farmers sow early, before the onset of the annual rains. Traditionally, seeds are deep planted (10 - 25 cm) to benefit from residual soil humidity and to protect from damage from late frosts. This practice allows varieties that require 160-180 days to reach maturity to be grown in areas with a frost-free season of 90-120 days. The volcanic soils of the Mexican highlands have low pH, restricting the availability of phosphate to the plant (Bayuelo-Jiménez *et al.*, 2011). Although displaying enhanced phosphorus use efficiency (Bayuelo-Jiménez & Ochoa-Cadavid, 2014), Mexican highland landraces tend to show restricted root growth (CIMMYT & Bjarnason, 1994; Eagles & Lothrop, 1994). To compensate for weak root development and prevent lodging, plants may be hilled (piling of soil around the base of the plant) up to three times during vegetative growth.

Palomero Toluqueño (PT) is a popcorn distributed in the highlands of the Mexican Central Plateau, notably in the valley of Toluca, at elevations from ∼2100 to ∼2900 m.a.s.l. (Fig. 1A. (Wellhausen *et al.*, 1951; Ruiz Corral *et al.*, 2008; Perales & Golicher, 2014)). Although present day cultivation is limited (https://www.biodiversidad.gob.mx/diversidad/proyectoMaices), PT is considered ancestral to the broader Mexican highland maize group and a progenitor of more productive modern highland landraces (Reif *et al.*, 2006; Arteaga *et al.*, 2016). PT has a relatively small genome and was selected as the target of the first landrace maize genome sequencing study (Vega-Arreguín *et al.*, 2009; Vielle-Calzada *et al.*, 2009). Subsequent work has continued to explore gene expression variation in PT (Aguilar-Rangel *et al.*, 2017; Crow *et al.*, 2020) and characterize wild-relative introgression through generation of additional genomic sequence (Gonzalez-Segovia *et al.*, 2019). In contrast, B73 is an elite inbred line developed in 1972 by Iowa State University as part of the breeding program for the US maize corn belt. It was highly prized for its ability to form high yielding hybrids and soon became a key ancestral female line in today’s global germplasm pool (Troyer, 1999; Iowa State University, 2009). B73 was selected for the first genome assembly in maize (Schnable *et al.*, 2009) and remains as the primary reference genome (Jiao *et al.*, 2017).

**Figure 1.**
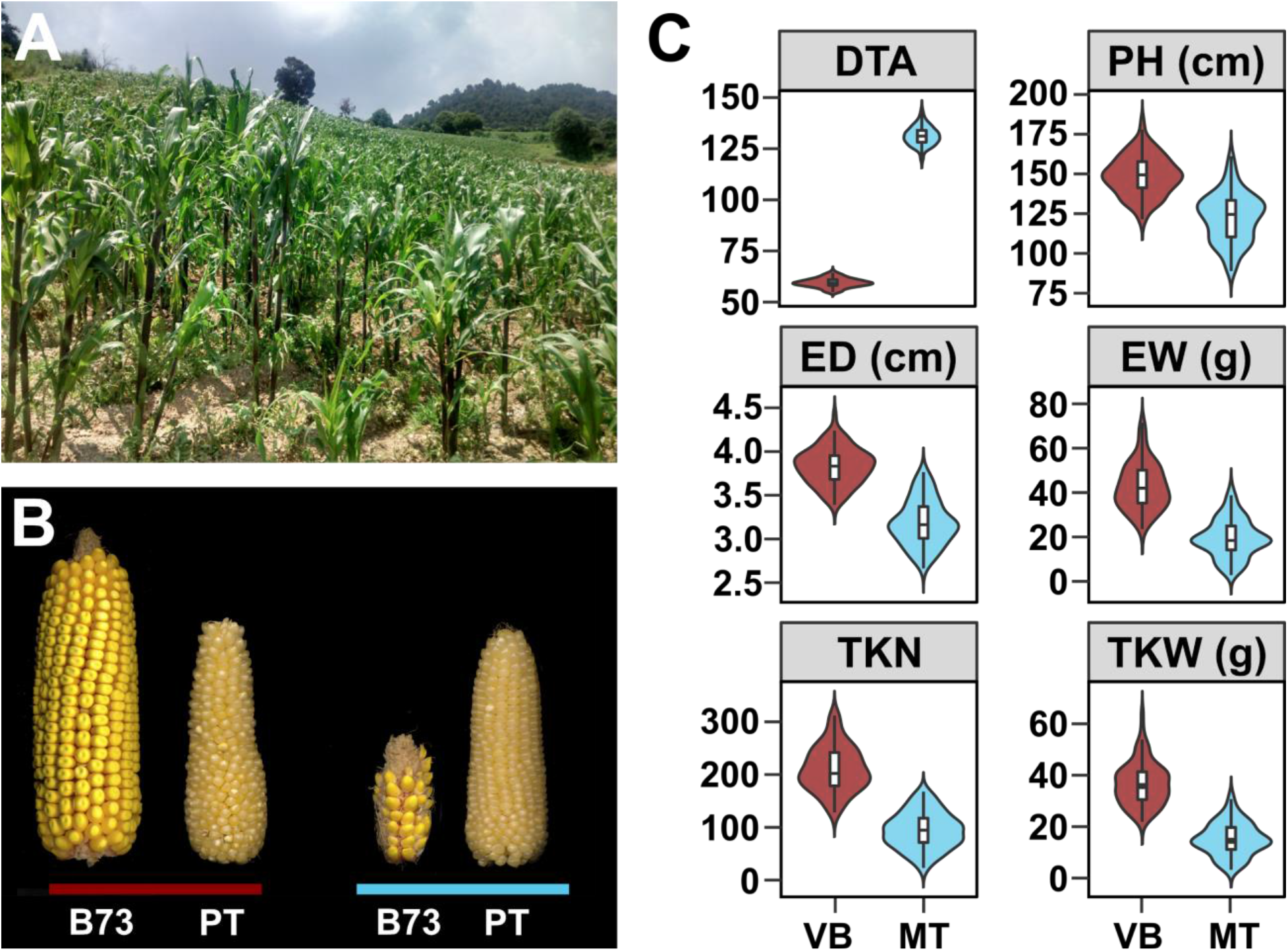
The highland environment impacts maize growth and productivity. A) A representative highland cultivated maize field at 3000 m.a.s.l. near the Nevado de Toluca volcano, State of Mexico, Mexico (19.121702, - 99.660812). B) Representative ears of the US adapted inbred line B73 and the Mexican highland landrace Palomero Toluqueño (PT) grown at sea level (red bar; 54 m.a.s.l.; Valle de Banderas [VB], Nayarit, Mexico) or in the highlands (blue bar; 2610 m.a.s.l.; Metepec [MT], State of Mexico, Mexico). C) Effect of the highland environment on plant performance. Distribution of trait values for 122 B73xPT BC_1_S_5_ lines grown in the lowland (VB; red) and highland (MT; blue) field sites. Trait codes: DTA - days to anthesis (days); PH - plant height (cm); ED - ear diameter (cm); EW - ear dry weight (g); TKN - total kernel number; TKW - total kernel weight (g). Fitted values for each genotype and location combination were used to generate violin plots and were estimated by adding BLUPs of the G+GEI to the estimated location mean. Boxes represent the interquartile range with the horizontal line representing the median and whiskers representing 1.5 times the interquartile ranges. The shape of the violin plot represents probability density of data at different values along the y-axis.

In this work, we characterize a mapping population generated from the cross of B73 and PT. We demonstrate local adaptation in PT and characterize associated genetic architecture by reciprocal transplant and two-site evaluation of our mapping population. We identified Quantitative Trait Loci (QTL) linked to phenology, morphology and yield components, including evidence to QTL x environment interaction (QEI). Overall, morphological QTL were stable across environments with either little QEI or mild scaling effects. We observed stronger QEI associated with yield components, including an example of antagonistic pleiotropy on chromosome (chr) 7, indicating a single locus fitness trade-off. We found evidence for relatively complex genetic architectures associated with putatively adaptive morphological traits. We discuss the implications of these results with respect to the origin of adaptive variation during rapid local adaptation in cultivated species.

## MATERIALS AND METHODS

### Plant material

To generate the biparental mapping population, an F_1_ was generated from the cross between the reference inbred line B73 and pollen pooled from several individuals of Palomero Toluqueño (PT), an open pollinated landrace endemic of the Mexican highlands. The accession used was MEXI5 (CIMMYTMA-002233) obtained from the International Center for Maize and Wheat Improvement (CIMMyT) seed bank, originally collected near the city of Toluca, Mexico State (19.286184N,−99.570871W) at 2597 m.a.s.l. A single B73xPT F_1_ individual was crossed as male to multiple B73 ears to generate a large BC_1_ population, capturing a single haplotype of PT. The BC_1_ was then self-pollinated 5 generations to form a BC_1_S_5_ RIL population, with an average of 25% of their genome from PT, and 75% of B73. 120 different families were advanced as independent pedigrees from BC_1_S_1_ to BC_1_S_5_. The same initial crossing strategy was used to generate material from the cross between B73 and the open-pollinated Conico/Celaya accession Michoacán 21 (Mi21; CIMMYTMA-001872). B73xMi21 stocks were further backcrossed to B73 with phenotypic selection for sheath pubescence and a segregating BC_5_S_1_ stock produced. The progenitor B73xPT and B73xMi21 F_1_ individuals described here are the same as those used in a previous report to derive introgression stocks segregating the *Inv4m* inversion polymorphism (Crow *et al.*, 2020).

### DNA preparation and genotyping

DNA was extracted from 100 B73xPT Recombinant Inbred Lines and four different B73 and PT individuals using isopropanol extraction. 50 mg of leaf tissue were harvested for each plant in a 2.0 mL tube and then frozen to −80 C. The frozen tissue was ground in a Qiagen TissueLyser II (Cat. ID: 85300) with a 30 Hz frequency for 30 s. After grinding, 300 μL of UEB1 (250 mM NaCl, 200 mM Tris pH 7.5, 25 mM EDTA, 0.5%SDS) buffer were added and the solution was mixed in a Thermomixer at 38 C for 10 minutes. 2μL of PureLink RNAse were added and the mix was left incubating for 30 min. After incubation, samples were separated by centrifugation at 14,000 rpm for 10 min at room temperature. 250 μL of supernatant was recovered and collected in a 1.5 mL tube. 40 μL of 3 M sodium acetate, pH 5.2 and 450 μL of isopropanol were added per tube, and samples incubated for 20 min at 4 C. A further centrifugation step was performed (14000 rpm, 10 min, room temperature) and the supernatant was discarded. Pellets were washed twice with 250 μL of 70% ethanol. The supernatant was discarded, and the pellet was left to dry for 30 min. When the pellet was dry, it was resuspended in 100 μL of milliQ water. DNA was quantified by spectroscopy and adjusted to a concentration of 20 ng/μL. DNA was genotyped at the SAGA (Servicio de Análisis Genético para la Agricultura, https://seedsofdiscovery.org/about/genotyping-platform/) laboratory in CIMMyT by DArTSeq (Edet *et al.*, 2018), generating ∼ 30,000 short length reads per sample.

### Processing of short read genotyping data and construction of the genetic map

Short read DNA sequences generated DArT-Seq were aligned to the v4 B73 reference genome (Jiao *et al.*, 2017) using seqmap (Jiang & Wong, 2008). Sequences that aligned to more than one physical position in the reference genome or that did not align were discarded. Genotype and SNP calling were performed with TASSEL 5 (Bradbury *et al.*, 2007). SNP calls were transformed to an ABH format, A assigned to B73 and B to PT. Sites for which the parental genotype was missing, ambiguous or heterozygous were removed. SNP calls were processed using Genotype-Corrector (Miao *et al.*, 2018), which considerably increased the contiguity of haplotypes among chromosomes. A set of 2, 067 polymorphic markers were selected for further analysis. The ABH genotype file was visualized using R/ABHgenotypeR (Reuscher & Furuta, 2016). Linked markers with shared patterns of segregation were identified with findDupMarkers function of R/qtl package (Broman *et al.*, 2003). Removing redundant makers reduced the final set to 918 polymorphic markers. The linkage map was built using the R/ASmap::mstmap (Taylor & Butler, 2017) under the Kosambi map function. Five individuals from a B73xMi21 BC_5_S_1_ family segregating sheath pubescence were genotyped using the same DArT-Seq platform as part of a project described in (Gonzalez-Segovia *et al.*, 2019).

### Field Evaluation

The BC_1_S_5_ population was evaluated in the highlands during 2015, 2016, 2018 and 2019 at 2610 m.a.s.l. in Metepec (MT; Mean average temperature: 12.4 °C; mean annual precipitation: 809 mm; Andosolic soil), Mexico State, and in the lowlands during 2015 and 2016 at 54 m.a.s.l. in Valle de Banderas (VB. Mean average temperature: 25.8 °C; mean annual precipitation: 1173 mm; Regosolic soil), Nayarit (Table S1; Fig. S6). BC_1_S_5_ families were evaluated in single-row 15 plant plots with 15 plants in 3 randomized complete blocks in MT and two blocks in VB. B73 and PT parents were inserted randomly in each block during 2015 and 2016. Weeds and insects were controlled by chemical methods as needed. The VB field site was provided with a ferti-irrigation. System. The MT field site was rain-fed with supplemental sprinkler irrigation after planting and when needed.

### Data preparation and trait estimation

Preparation of trait data and QTL mapping was performed in R Statistics (Robinson *et al.*, 2010; McCarthy *et al.*, 2012; R Core Team, 2019). Data collected from single row plots were collapsed to a single value per plot: plot medians were taken for traits scored on multiple individuals; plot level traits such as stand count or flowering time were unchanged. Data was trimmed to remove outliers per trait/location (VB or MT) using R/graphics::boxplot default criteria. Continuous traits (ASI, DTA, DTS, ED, EH, EL, EW, PH, TKN, TKW, and TL) were further adjusted on a per block basis to a spline fitted using R/stats::smooth.spline against row number to reduce spatial variation at the sub-block scale. Spline fitting was not applied to any block containing less than 50 plots. The final dataset contained 4 years of data for location MT (123 genotypes from one block in 2015, 105 genotypes from three blocks in 2016, 140 genotypes from two blocks in 2018, and 110 genotypes from one block in 2019), and two years of data for location VB (123 genotypes from one block in 2015, and 117 genotypes from two blocks in 2016). For each continuous phenotypic trait, a mixed linear model was fitted using restricted maximum-likelihood with R/lme4::lmer. To fit the model, a location-year variable was generated to represent the location by year combinations, and a location-year-block variable was generated to represent all location, year, and block combinations, such that:

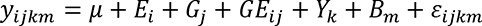

where the response variable *y*_*ijkm*_ is a function of the overall mean (*μ*), fixed effect of location (*E*_*i*_), random effect of genotype (*G*_*j*_), genotype by location interaction (*GE*_*ij*_), location-year term (*Y*_*k*_), location-year-block term (*B*_*m*_), and the residual. BLUP values for the genotypic effect (G) and genotype by location interactions (GEI) were extracted using R/lme4::ranef. We calculated BLUP values for each genotype and location combination (G+GEI) by adding genotypic BLUPs and GEI BLUPs (Olivoto *et al.*, 2019). We also calculated fitted values by adding BLUPs to the appropriate means for data visualization and downstream analyses using natural units: for genotypic effect, fitted values were calculated by adding genotypic BLUPs to the grand mean; fitted values for each genotype and location combination were calculated by adding G+GEI BLUPs to the location mean.The significance of the environment effect was evaluated by comparing the full model with location effect and the reduced model without location effect using the likelihood ratio test for continuous phenotypic traits (Table 1). For phenotypic traits with count and scale data, two-group Wilcoxon tests were conducted to evaluate the difference between the two locations (Table 1).

**Table 1:** Description of phenotypic traits measured at two locations. The significance of location effect was calculated using a Wilcoxon test for count and scale data (i.e., TBN, KPR, KRN, P_INT and Hair_score), and likelihood ratio test was conducted for the other continuous traits. Note: *: p < 0.05; **: p < 0.001; ***: p < 0.0001.

| Phenotypic traits | Description | Unit | Mean (MT) | Mean (VB) | Significance of E |
| --- | --- | --- | --- | --- | --- |
| DTA | Days to anthesis | Day | 131.07 | 59.69 | < 0.0001*** |
| DTS | Days to silking | Day | 131.30 | 60.15 | < 0.0001*** |
| ASI | Anthesis-silking interval | Day | 0.46 | 0.30 | 0.5650 |
| PH | Plant height | cm | 122.72 | 148.65 | 0.0546 |
| TBN | Tassel branch number | Count | 4 | 5 | 0.1796 |
| EW | Ear weight | g | 19.60 | 42.69 | 0.0019** |
| EL | Ear length | cm | 7.30 | 8.75 | 0.0860 |
| ED | Ear diameter | cm | 3.18 | 3.83 | 0.0083** |
| EH | Ear height | cm | 49.68 | 65.30 | 0.0260* |
| KRN | Number of kernel rows | Count | 14 | 17 | 0.0157* |
| KPR | Number of kernels per row | Count | 13 | 20 | 0.0275* |
| TKW | Total kernel weight | g | 15.31 | 36.50 | 0.0002** |
| TKN | Total kernel number | Count | 94.44 | 210.14 | 0.0001** |
| TL | Tassel length | mm | 21.65 | 28.35 | 0.0573 |
| P_INT | Anthocyanin pigment intensity | visual 0-4 code scale | 2 | 0 | 0.5303 |
| MH | Macrohair score | Hair pattern<br><= 1, hair score =0;<br>Hair pattern >1, hair score = hair density | 0 | 0 | --- |

### QTL mapping

The BLUPs for continuous traits, the medians for count traits and the mode for semi-quantitative scale traits were used as phenotypic inputs for QTL mapping. Phenotypic scores were selected/combined to perform four distinct analyses: 1) GEN: the genotype main effect (G) of the mixed linear model for continuous traits and the median/mode across all plots for other traits; 2) VB: G+GxE term for VB for continuous traits and the VB median/mode for other traits; 3) MT: G+GxE term for MT for continuous traits and the MT median/mode for other traits; 4) GEI: the difference between MT and VB GxE BLUPs for continuous traits and the difference between MT and VB median/mode for other traits.

Individual QTLs were detected using single QTL scan and Multiple QTL Mapping (MQM) with R/qtl::scanone (default options; Haley-Knott regression. Broman *et al.*, 2003) and R/qtl::MQM (default options. 100 autocofactors, step.size = 1, window.size = 25. Arends *et al.*, 2010), respectively. Genome-wide LOD significance thresholds were established at α = 0.05 by 1000 permutations of scanone and MQM and scanone models. Individually QTL were combined in an additive multi-QTL model with R/qtl::makeqtl and their positions refined with R/qtl::refineqtl. The function R/qtl::addqtl was used to detect additional QTLs in a multi-QTL context with a LOD threshold of 3 LOD considered significant. The final multi-QTL model was applied using R/qtl::fitqtl (Haley-Knott regression) to obtain the refined position and variance explained. Significance levels of the full model and the component QTL terms were obtained from the drop-one ANOVA table. Bayes confidence intervals were obtained from the fitqtl model. The effect size and effect plots of each individual term of the full model were obtained with R/qtl::effectplot.

### Elevation eGWAS

We performed an environmental genome-wide association analysis (eGWAS) to measure the association between genetic loci and the elevation of native environment for landrace accessions across Mexico, as previously described (Gates *et al.*, 2019). The data set consisted of 1, 830 Mexican maize landrace accessions from the CIMMyT Maize Germplasm Bank with elevation data, genotyped for 440, 000 single nucleotide polymorphisms (SNPs; Romero Navarro *et al.*, 2017; Gates *et al.*, 2019). We used a linear model to fit the genotypic data and elevation to the landrace data. The first five eigenvectors of the genetic relationship matrix were included in the linear model to control for the population structure. The top 1000 SNPs with the strongest association with elevation were selected and used in the downstream analysis.

### Data availability statement

PT (CIMMYTMA-002233) and Mi21 (CIMMYTMA-001872) accessions are available directly from CIMMyT (www.cimmyt.org). All other materials are available on request subject to costs of propagation and export if outside of Mexico. Derived material is covered by the same CIMMyT MTA as the progenitor parents. Supplemental files available at https://doi.org/10.6084/m9.figshare.16608517.v1. *Phenotypic data*: BLUPs and fitted values estimated for diverse traits for 97 B73xPT BC_1_S_5_ families. *Genetic map*: genetic map of the B73xPT BC_1_S_5_ mapping population. *Altitude* eGWAS: results contains the results of the eGWAS analysis. *QTL LOD profile*: LOD profile of the multi-QTL models for different traits for each set of phenotypic data. *Effect Plots*: estimated effect of the QTLs detected for a set of phenotypic data using fitted values. *Reaction norms*: contains the reaction norms estimated for all the QTLs detected in VB and MT phenotypic sets using fitted values.

## RESULTS

### The stress of the highland environment limits maize growth and productivity

To characterize the genetic architecture of highland adaptation in Mexican native maize (Fig. 1A), we crossed the highland landrace Palomero Toluqueño (PT) to the US reference inbred line B73 and derived 120 BC_1_S_5_ families. When generating the BC_1_, we used a single F_1_ individual as a male to pollinate several B73 females, ensuring that a single PT haplotype was captured from the open-pollinated donor accession. As a consequence our mapping population was bi-alleleic, *i.e.* segregating for B73 and a single PT allele at any given locus. The final BC_1_S_5_ families carried ∼25% PT genome in a B73 background, with homozygosity > 98%. BC_1_S_5_ families were genotyped using the DArT Seq platform (http://www.diversityarrays.com/) and a final set of 918 markers were selected and used to generate a linkage map for quantitative trait locus (QTL) mapping.

We evaluated the 120 B73 x PT BC_1_S_5_ families and parents in Mexican lowland (Valle de Banderas, Nayarit at 54 m.a.s.l.) and highland (Metepec, Mexico State, at 2610 m.a.s.l.) field sites. Lowland trials were conducted during the dry season from November to March with supplemental irrigation. Highland trials were conducted in a rain-fed field in the standard highland cycle from April to November. We collected data on a range of phenological, morphological and agronomic traits (Table 1). B73 and PT parents showed a classic pattern of rank-changing GEI for yield components across the two locations, demonstrating adaptation of PT to the highland environment (Fig. 1B, 1-S5). The negative impact of the highland environment on B73 was dramatic, while PT was more stable across the two sites. Across all of the BC_1_S_5_ genotypes, there was a significant environmental effect on 9 of 19 traits (Fig. 1C; 1-S1; Table 1). Average flowering (days to anthesis, DTA and days to silking, DTS), measured in chronological days, was greatly delayed in the highland site by 71 days. Overall, plants in the highlands were shorter in stature (plant height, PH) and produced smaller ears (ear diameter, ED; ear height, EH; ear weight, EW) bearing fewer grains (total kernel number, TKN) (Fig. 1C). Average total kernel weight per plant (TKW) dropped from 36.5 g to 15.3 g, a reduction of 58 %, from the lowland to highland field (Fig.1 C).

### Segregation in the BC_1_S_5_ reflects GEI seen in the B73 and PT parents

Having characterized the main effect of the highland environment, we explored GEI among the BC_1_S_5_ families. For certain traits, such as EH, there was a clear environmental effect (Fig. 2A), but little evidence of GEI between the parents nor among the lines (Fig. 2B). In contrast, yield components such as EW and TKW showed a strong environmental effect (Fig. 1C), GEI between parents, and extensive rank-changing GEI among BC_1_S_5_ families (Fig. 2C, D; Fig. 2-S1). Taking EW as a proxy for yield, many BC_1_S_5_ families were more stable with respect to location than the two parents, although the majority were inferior to the better parent in either site. That said, we did observe a small number of families that performed as well as, or better, than the parents in both locations.

**Figure 2.**
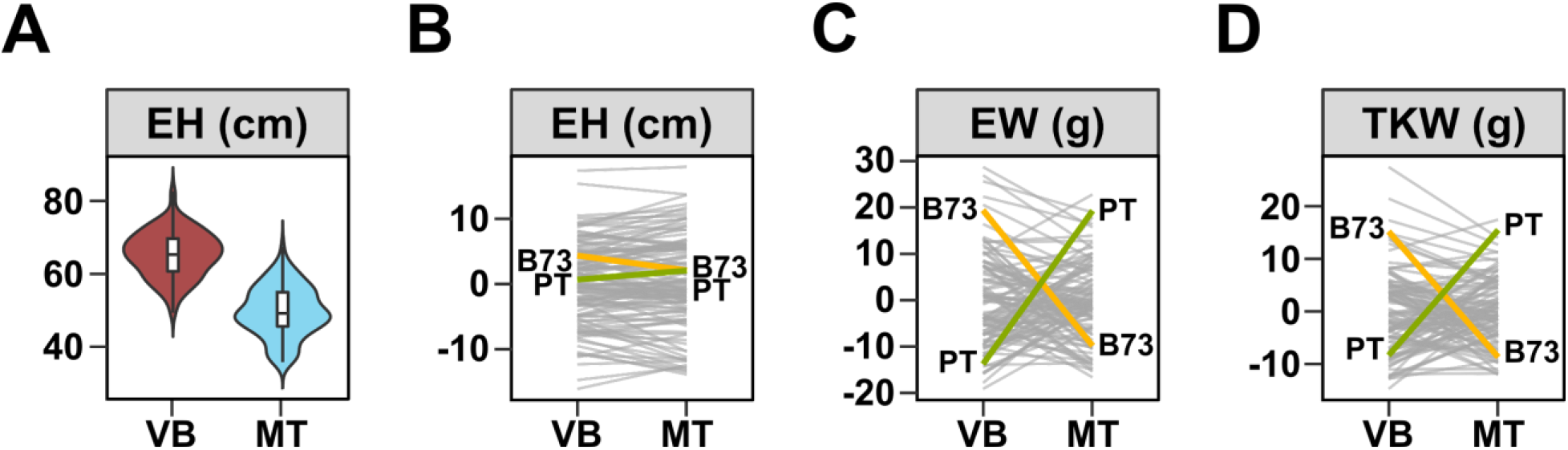
Extensive GEI was observed for yield components. A) Distribution of *ear height* (EH, cm) in low (VB) and high elevation (MT) field sites. Boxes represent the interquartile range with the horizontal line representing the median and whiskers representing 1.5 times the interquartile range. The shape of the violin plot represents probability density of data at different values along the y-axis. B) Reaction norm plot for EH, showing little GEI. Values shown are G + GEI deviations from the field site average. Line segments connect values for each RIL genotype in the two field sites. B73 (yellow) and PT (green) parental values are shown. C), D) as B, showing extensive rank-changing GEI associated with *ear weight* (EW, g) and *total kernel weight* (TKW, g), respectively.

### QEI associated with yield components indicates local adaptation at the locus level

Extensive GEI for yield components could be associated with either conditional effects or antagonistic pleiotropic at individual QTL (Fig. 3A). To explore the genetic architecture underlying GEI for yield components in our BC_1_S_5_ families, we performed a QTL analysis, using a series of trait combinations to allow us to determine QTL effects in the lowland and highland sites, as well as to identify QEI. Across the four phenotypic sets, we identified 44 different QTLs where eighteen were significant under the GxE analysis, mirroring the GEI observed at the level of individual lines (Table 2).

**Figure 3.**
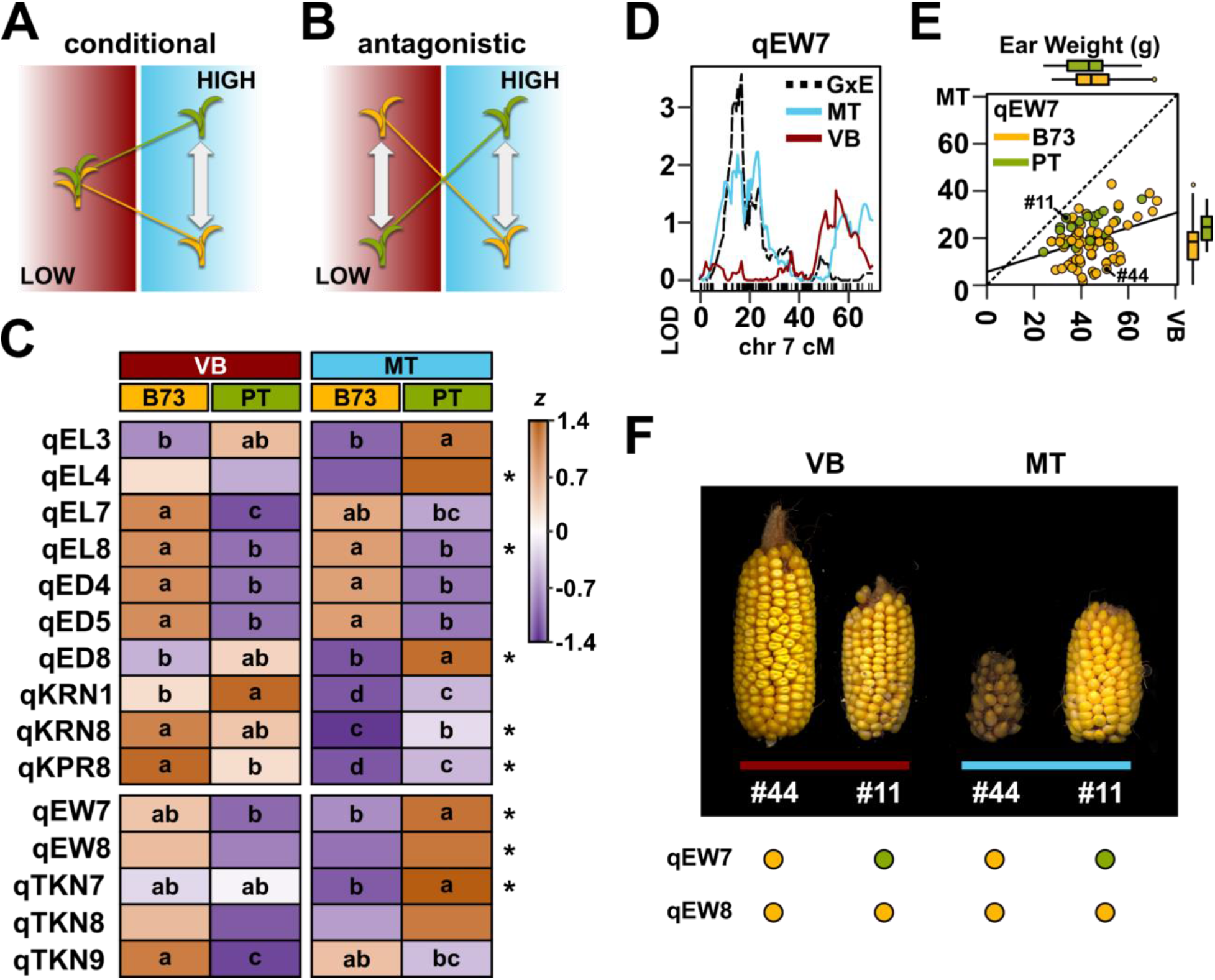
QEI interactions contribute to local adaptation. A,B) Schematic of QEI under models of A) *conditional* effect in one environment but not another or B) *antagonistic pleiotropy* showing a change in the sign of the QTL effect between environments. C) Heatmap representation of the standardized median G + GEI value for all families with a given genotype (B73 or PT) in lowland (VB) and highland (MT) sites, for the named QTL (see Table 2). Asterisks indicate QTL identified in the GEI model. Lowercase letters indicate Tukey means groups. D) LOD support for a conditional *ear weight* (EW) QTL (qEW7) on the short arm of chromosome (chr) 7. The QTL is well supported by data from the highland site (MT, blue trace) but not the lowland site (VB, red trace), and is captured by a mutiQTL model for GxE (black trace). E) Scatter plot of EW in highland (MT) against lowland (VB) fields. Each RIL is represented by a single point, colored by genotype at qEW7 (yellow, B73; green, PT).RILs shown in F below are labelled. The solid line shows a linear fit through all points. Box plots parallel to the vertical and horizontal axes show the distribution by genotype in MT and VB, respectively. Boxes represent the interquartile range with the horizontal line representing the median, and whiskers extending 1.5 times the interquartile range. F) Ears of RILs LANMLR17B044 (#44) and LANMLR17B011 (#11) produced in lowland (red bar) and highland (blue bar) fields, showing marked differences in stability with respect to field. Points below the panel indicate QTL genotype.

**Table 2:** QTLs detected in the B73 x PT BC_1_S_5_ RIL population for the phenotypes (Trait, abbreviated as in Table 1) and datasets (Set Detected) analyzed. Marker column describes the marker linked to the QTL, chr: chromosome; Pos: genetic position of the QTL LOD peak (cM); P_pos: physical position of the QTL LOD peak (MB, Reference genome v4), SI: Support Interval of the QTL (MB); GxE: indicates if the QTL is detected in the GxE data set; %VE: additive variance explained by the QTL in the multi-QTL model.

| QTL | Marker | Chr | pos | p_pos | Set Detected | SI | GxE | % VE |
| --- | --- | --- | --- | --- | --- | --- | --- | --- |
| qASI1 | 1_53029437 | 1 | 30.46 | 53.03 | VB | 35.2 - 292.18 |  | 10.13 |
| qASI2 | 2_241675850 | 2 | 77.91 | 241.68 | GEN, VB | 240.52 - 244.41 |  | 10.88 - 14.54 |
| qASI3 | 3_229390098 | 3 | 65.63 | 229.39 | GEN | 11.85 - 234.87 |  | 7.29 |
| qASI8 | 8_135484500 | 8 | 24 | 135.48 | GEN, VB, MT | 134.32 - 164.79 |  | 15.96 - 18 |
| qDTA1 | 1_286172395 | 1 | 94.35 | 286.17 | VB, GxE | 2.81 - 306.46 | * | 6.07 - 10.19 |
| qDTA6 | 6_166664744 | 6 | 22 | 166.66 | GEN, MT, GxE | 165.39 - 168.82 | * | 7.5 - 13.53 |
| qDTA7 | 7_165928844 | 7 | 48.37 | 165.93 | MT | 0.84 - 173.75 |  | 5.51 |
| qDTA8a | 8_21391040 | 8 | 0 | 21.39 | MT | 25.12 - 97.57 |  | 9.45 |
| qDTA8b | 8_153580487 | 8 | 27.5 | 153.58 | GEN, VB, MT, GxE | 148.95 - 161.29 | * | 9.67 - 21.23 |
| qDTS1 | 1_12171293 | 1 | 12.47 | 12.17 | GxE | 2.81 - 306.46 | * | 5.18 |
| qDTS6 | 6_166506716 | 6 | 23 | 166.51 | GEN, MT, GxE | 165.39 - 168.12 | * | 7.59 - 15.18 |
| qDTS7 | 7_163657186 | 7 | 48 | 163.66 | MT | 15.91 - 169.56 |  | 8.16 |
| qDTS8 | 8_161289005 | 8 | 31 | 161.29 | GEN, MT, GxE | 133.04 - 169.05 | * | 11.17 - 16.07 |
| qED4 | 4_179539232 | 4 | 41.65 | 179.54 | GEN, VB | 163.36 - 196.26 |  | 14.69 - 14.96 |
| qED5 | 5_190812247 | 5 | 31.54 | 190.81 | VB | 1.48 - 198.95 |  | 9.2 |
| qED8 | 8_97570666 | 8 | 7.81 | 97.57 | GxE | 21.39 - 112.91 | * | 14.86 |
| qEH1 | 1_177987239 | 1 | 53.68 | 177.99 | GEN, MT | 162.88 - 198.3 |  | 11.87 - 14.99 |
| qEH7 | 7_135399817 | 7 | 33.26 | 135.4 | GEN, VB, MT | 131.23 - 144.37 |  | 14.94 - 16.76 |
| qEL3 | 3_190627575 | 3 | 43.33 | 190.63 | MT | 11.5 - 234.87 |  | 13.39 |
| qEL4 | 4_74666833 | 4 | 23.02 | 74.67 | GxE | 18.12 - 161.42 | * | 12.34 |
| qEL7 | 7_172341845 | 7 | 55 | 172.34 | VB | 166.94 - 181.12 |  | 12.16 |
| qEL8 | 8_175513459 | 8 | 43.5 | 175.51 | GEN, VB, GxE | 172.04 - 179.51 | * | 9.25 - 14.81 |
| qEW7 | 7_20110508 | 7 | 16.9 | 20.11 | GxE | 9.14 - 28.5 | * | 14.33 |
| qEW8 | 8_110436593 | 8 | 13.45 | 110.44 | GxE | 88.36 - 170.14 | * | 11.6 |
| qMH3 | 3_156124621 | 3 | 28.83 | 156.12 | GEN, VB | 11.85 - 161.43 |  | 9.43 - 11.07 |
| qMH7 | 7_155328976 | 7 | 43.1 | 155.33 | GEN, MT | 152.38 - 164.52 |  | 14.21 - 17.8 |
| qMH8 | 8_122414801 | 8 | 18.71 | 122.41 | VB, GxE | 115.01 - 123.81 | * | 15.42 - 17.91 |
| qMH9 | 9_65716548 | 9 | 12.5 | 65.72 | GEN, VB | 20.72 - 124.18 |  | 16.61 - 19 |
| qKPR8 | 8_135484500 | 8 | 24 | 135.48 | GxE | 115.01 - 164.79 | * | 18.86 |
| qKRN1 | 1_161556632 | 1 | 49.5 | 161.56 | VB | 8.68 - 296.87 |  | 12.63 |
| qKRN8 | 8_133038186 | 8 | 23.42 | 133.04 | GxE | 82.76 - 164.79 | * | 16.39 |
| qP_INT2 | 2_19456739 | 2 | 25.61 | 19.46 | GEN, VB, MT | 18.31 - 25.87 |  | 22.12 - 40.03 |
| qP_INT10 | 10_11796399<br>5 | 10 | 19.49 | 117.96 | GEN, VB, MT | 116.16 - 138.71 |  | 9.47 - 13.37 |
| qPH1a | 1_197051908 | 1 | 56.5 | 197.05 | GEN, VB, MT | 176 - 199.71 |  | 18.88 - 20.43 |
| qPH1b | 1_293830327 | 1 | 100.1<br>3 | 293.83 | VB | 283.08 - 299.44 |  | 12.26 |
| qPH8 | 8_149541560 | 8 | 27.58 | 149.54 | VB | 142.92 - 164.79 |  | 13.69 |
| qPH10 | 10_13688363<br>9 | 10 | 26 | 136.88 | GxE | 10.27 - 143.76 | * | 11.17 |
| qTBN2 | 2_141142660 | 2 | 40.88 | 141.14 | GEN | 70.9 - 184.93 |  | 13.24 |
| qTBN7 | 7_121548061 | 7 | 24.52 | 121.55 | GEN, VB, GxE | 112.71 - 121.69 | * | 15.86 - 33.17 |
| qTKN7 | 7_15912522 | 7 | 15.35 | 15.91 | MT | 9.14 - 121.69 |  | 17.09 |
| qTKN8 | 8_126886782 | 8 | 20.81 | 126.89 | GxE | 82.76 - 170.04 | * | 12.86 |
| qTKN9 | 9_124180939 | 9 | 19.6 | 124.18 | VB | 111.77 - 135.89 |  | 16.39 |
| qTL1 | 1_227715124 | 1 | 72.51 | 227.72 | GxE | 224.77 - 244.86 | * | 19.73 |
| qTL2 | 2_230998297 | 2 | 70.19 | 231 | GEN, MT | 0.94 - 241.68 |  | 11.56 - 11.62 |

We identified a total of 14 QTL for the yield components: ear length (qEL3, qEL4, qEL7, and qEL8), ear diameter (qED4, qED5, and qED8), kernel row number (qKRN1 and qKRN8), kernels per row (qKPR8), ear weight (qEW7 and qEW8), and total kernel number (qTKN7, qTKN8, and qTKN9) (Fig. 3C, Table 2). QTL for the primary yield component EW on chromosomes (chr) 7 and 8 showed the B73 alleles to be associated with higher EW in the lowlands, and the PT alleles associated with higher EW in the highlands (Fig. 3C). Statistical support for GEI was detected at EW7 but not EW8. There was some support for antagonistic pleiotropy at EW7, although evidence was strongest for a highland conditional effect (Fig. 3D, E, F).

### PT alleles at major flowering time QTL on chromosome 8 and 6 accelerate flowering

We detected 13 unique QTLs for flowering time traits (DTA, DTS, ASI) some of which were detected across all analyses (*e.g.* qDTA8b, detected in GEN, VB, MT and GEI sets). Clusters of flowering QTL were found on both chromosomes 8 and 6 (Table 2). qDTA8b, qDTS8 and qASI8 were consistently detected across the GEN, MT, VB and GEI sets, and qDTA6, qDTS6 were detected in the GEN, MT and GEI sets. Mirroring the parental difference, the PT allele at qDTA8b, qDTS8, qDTA6 and qDTS6 accelerates flowering. Flowering QTL have been consistently detected on Chr 8 in maize-teosinte mapping populations and maize diversity panels in the context of differences between temperate and tropical material (Jiang *et al.*, 1999; Chardon *et al.*, 2004; Buckler *et al.*, 2009; Coles *et al.*, 2010; Bouchet *et al.*, 2013; Xu *et al.*, 2017; Guo *et al.*, 2018). The QTLs on chromosome 8 are in the vicinity of the well-characterized flowering loci *vgt1* and *ZEA CENTRORADIALIS 8* (*Zcn8*). The locus *vgt1* corresponds to a non-coding region of ∼ 2 kb that regulates *ZmRap2.7*, an *APETALA-2* like gene located ∼ 70 kb downstream (Salvi *et al.*, 2007); *Zcn8* is the florigen gene of maize and has a central role in mediating flowering (Meng *et al.*, 2011; Guo *et al.*, 2018). Polymorphisms in *vgt1* and *Zcn8* have previously been associated with flowering time variation associated with both adaptation to latitude and altitude (Salvi *et al.*, 2007; Ducrocq *et al.*, 2008; Buckler *et al.*, 2009; Romero Navarro *et al.*, 2017; Guo *et al.*, 2018). Given their close proximity, it was not possible to confidently separate the potential effects of *vgt8* and *Zcn8* in our population, and we consider it possible that the combined effect of variation in these two loci underlies our Chr 8 QTL.

The QTL qDTA6 and qDTS6 are in close proximity to the gene *Peamt2* (Zm00001eb294690, chromosome 6 ∼ 166.5 MB), an ortholog of the *Arabidopsis XIPOTL1* gene which encodes a phosphoethanolamine N-methyltransferase (PEAMT). PEAMT catalyzes the transformation of phosphocholine to phosphatidylcholine (PC), (Cruz-Ramírez *et al.*, 2004; Sánchez Martínez, 2018). The balance between PC and its precursors has been associated with the timing of flowering in *Arabidopsis* (Nakamura *et al.*, 2014) and previously implicated in early flowering in Mexican highland maize (Rodríguez-Zapata et al., 2021). Nevertheless, fine mapping and metabolic studies would be needed to confirm the possible role of variation of *Peamt2* in flowering time variation

### Identification of QTL linked to characteristic tassel and stem traits

PT displays a number of putatively adaptive morphological traits that are characteristic of the Mexican highland group as a whole (Fig. 1A, Eagles & Lothrop, 1994; Gonzalez-Segovia *et al.*, 2019). To gain insight into the targets and mechanism of selection during local adaptation, we collected data on tassel (male inflorescence) morphology, stem pigmentation and stem pubescence from our BC_1_S_5_ families (Table 1).

The PT tassel is large but unbranched with respect to B73 or typical Mexican lowland landraces (Fig. 4A, B). Across the BC_1_S_5_ population, tassel length (TL) and tassel branch number (TBN) showed a mild reduction in the highlands compared with the lowland environment, with little GEI between parents or among families (Fig. 4C-F). We identified two QTL linked to TL (qTL1 and qTL2) and two linked to TBN ( qTBN2 and qTBN7; Table 2). In common with observations at the whole genotype level, QTL effects for tassel traits were constant in the two environments, and there was no indication of QEI <u>(Perez-Limon et al. 2021, QTL_reaction_norms)</u>. For qTL1, the effect was not aligned with the parental difference, with the PT allele being linked to shorter TL. For TBN, the PT allele at qTBN7 was linked to less branching, while the PT allele at qTBN2 was linked to greater branching (Figure 4H). The largest TBN effect was associated with qTBN7 that co-localized with a previously reported TBN QTL (Xu *et al.*, 2017; Gonzalez-Segovia *et al.*, 2019) and the *Ramosa1* (*Ra1*, Zm00001d020430, chromosome 7 at 113.57 MB) candidate gene (Sigmon & Vollbrecht, 2010; Fig. 3G, H).

**Figure 4.**
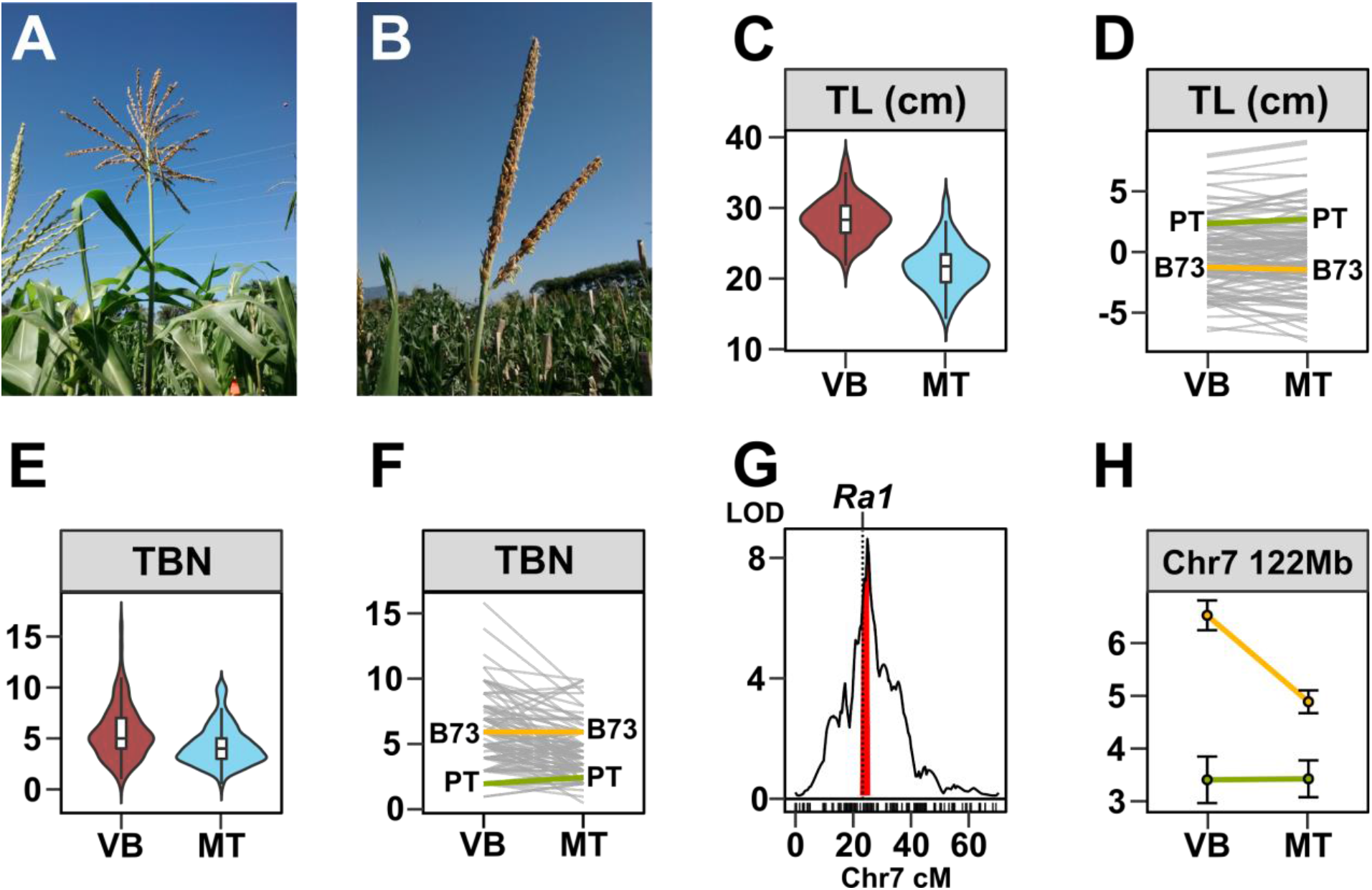
A major QTL for tassel branch number co-localizes with the *Ramosa1* gene. In comparison with typical maize varieties (A), tassel branching is strongly reduced in Mexican highland maize (B). C) Distribution of *tassel length* (TL, cm) in low (VB) and high (MT) field sites. Boxes represent the interquartile range with the horizontal line representing the median and whiskers representing 1.5 times the interquartile range. The shape of the violin plot represents probability density of data at different values along the y-axis. D) Reaction norm plot for TL. Values shown are G + GxE deviations from the field site average. Line segments connect values for each RIL genotype in the two field sites. B73 (yellow) and PT (green) parental values are shown. E, F) as C and D for *tassel branch number* (TBN). For F, the plot shows the median for each genotype in each field. G) LOD support (multQTL model, G main effect) for a qTBN7 that co-localizes with the *Ramosa1* (*Ra1*) candidate gene. Red shading indicates a drop 2 LOD interval around the peak marker. H) Effect of the chr 7 TBN QTL showing trait values for families carrying B73 (yellow) or PT (green) alleles in lowland (VB) or highland (MT) field sites.

PT displays strong stem pigmentation in comparison to the non-pigmented stem of B73 (Fig. 1A, 5A). We detected two QTL for pigment intensity (qINT2 and qINT10) with no evidence of QEI (<u>Perez-Limon et al. 2021, QTL_reaction_norms</u>; Table 2). The qINT2 interval colocalizes with a QTL previously reported in a Palomero Toluqueño x Reventador F_2_ mapping population (Gonzalez-Segovia *et al.*, 2019). Pigment QTL were linked to the well-characterized basic helix-loop-helix (bHLH) regulators of anthocyanin biosynthesis *b1* (Zm00001d000236 chr 2, 198.2 MB) and *r1* (Zm00001d026147, chr10 139.78 MB. Dooner & Kermicle, 1976; Radicella *et al.*, 1992; Selinger *et al.*, 1998; Selinger & Chandler, 1999, 2001; Chatham & Juvik, 2021). It has been demonstrated that functional variation at *b1* is driven by varying patterns of upstream transposon insertion, leading to differences in pigmentation patterns. For example, *B-Bolivia* induces the biosynthesis of anthocyanin in both vegetative tissue and the aleurone of the grain, while the *B-Mex7* allele, which was identified from the Mexican highland landrace Cacahuazintle, induces pigment in the sheath margins of the sheath (Chandler *et al.*, 1989; Radicella *et al.*, 1992; Selinger & Chandler, 1999).

**Figure 5.**
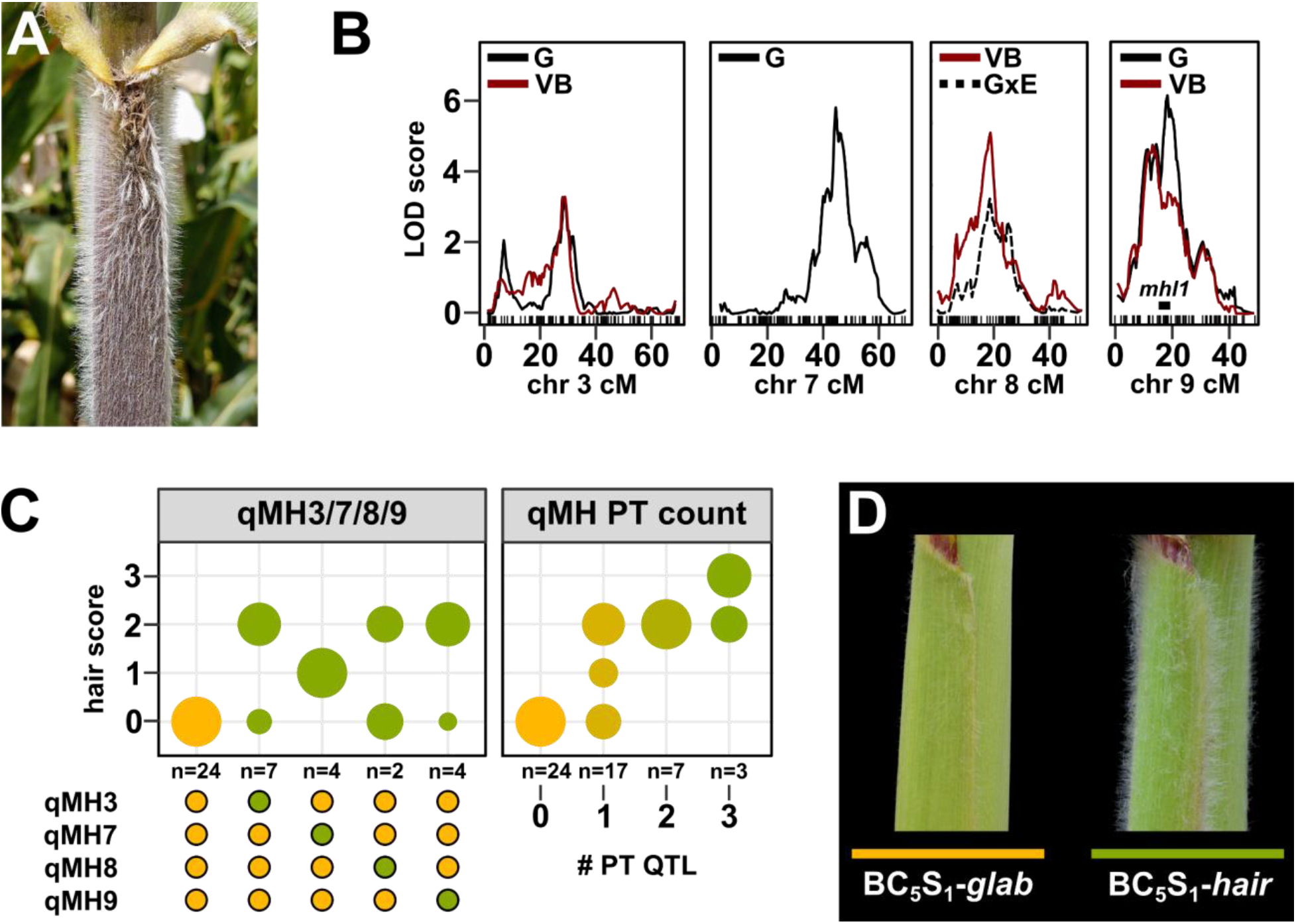
Stem macrohair production is promoted by multiple QTL. A) Mexican highland maize is characterized by extensive sheath pubescence. B) QTLs linked to macrohair score (MH) on chromosomes (chr) 3, 7, 8 and 9. Trace shows LOD support for the macrohair trait in the lowland field (VB), in the genotype main effect (G) or for GEI (GxE). Teosinte introgression on chr 9 reported by Hufford *et al*., 2013 that includes the *mhl1* locus is marked by a black bar. C) QTL effect shown as the proportion (shown by circle diameter in the main plot) of RILs scored for different hair score values in a given genotypic class (B73 allele, yellow; PT allele, green). Panels show the effect of allele substitution at the stated QTL in the subset of RILs for which the other QTLs are fixed as B73 and the cumulative effect of increasing the number of PT alleles at qMH 3, 7, 8 or 9. Points below the panel indicate QTL genotype. D) Glabrous (*glab*) and pubescent (*hair*) near-isogenic siblings generated by selection for pubescent plants through five generations of backcrossing of a Mexican Conico highland landrace to B73.

PT, in common with other Mexican highland landraces, exhibits pronounced stem pubescence (Fig. 5A). Although stem macrohairs were present in the BC_1_S_5_ families, no single family reached the level of pubescence seen in the PT parent, suggesting a complex genetic architecture. Furthermore, the reduced vigor of the BC_1_S_5_ families in the highland location was associated with poor expression of the pubscence trait and difficulty in scoring. Using a semi-quantitative scale for evaluation, we identified four QTL linked to stem macrohairs, located on chromosomes 3, 7, 8, and 9 (Fig. 5). The QTL interval qMH9 included the *macrohairless1* (*mhl1*, bin 9.04) locus that has previously been linked to the production of leaf blade macrohairs in temperate inbred maize (Moose *et al.*, 2004). The qMH9 region also coincided with a previously reported region of introgression from the highland teosinte *Zea mays* ssp. *mexicana* (itself typically pubescent) to Mexican highland maize (Hufford *et al.*, 2013; Gonzalez-Segovia *et al.*, 2019; Calfee *et al.*, 2021). This region has been characterized as a chromosomal inversion of ∼3 MB that displays patterns of selection in highland maize populations (Calfee *et al.*, 2021). The qMH9 interval was relatively large (∼12 cM, estimated to cover ∼100 Mb) and inspection of the LOD profile suggested the possible presence of two peaks (Fig. 5B). Although presented here as a single QTL, there may in fact be two linked factors.

For all macrohair QTL, the PT allele was associated with greater stem pubescence. We previously reported difficulty in mapping sheath macrohairs in a PT x lowland landrace F_2_ population because nearly all plants were scored as pubescent in a simple qualitative evaluation (Gonzalez-Segovia *et al.*, 2019). We interpreted this previous observation to indicate the action of several partially dominant factors, each individually sufficient to trigger the production of stem macrohairs. To further test this hypothesis, we extracted the effect of the PT allele at each macrohair QTL in turn, fixing the other loci as B73 (Fig. 5C). Consistent with genetic redundancy, the PT allele at any macrohair QTL was sufficient to promote a degree of stem pubescence (p < 0.01 for all three QTL compared to families carrying B73 alleles at all qMH loci). Although limited by the size of our population and the qualitative nature of our phenotyping, we could detect a significant difference between families carrying PT alleles at several macrohair loci and those carrying the PT allele at only one of the loci (p = 0.26; Fig. 5D). Unfortunately, no family carried PT alleles at all four of the loci (this is not unexpected in a BC_1_ population of 120 families). Although the four macrohair loci were individually sufficient to induce stem macrohair production, we hypothesise that their combined effect (and potentially that of additional loci) is necessary to approach the levels of pubescence of the PT parent.

In parallel with generation of the BC_1_S_5_ population, we also produced pubescent near isogenic lines (NILs) by phenotypic selection and recurrent backcrossing to B73. Here, we initially used several different Mexican highland landrace donors. Material generated from the PT relative Conico (accession Michoacan 21) consistently showed the greatest pubescence and was prioritized for backcrossing and genotypic analysis. A BC_5_S_1_ family showed 3:1 segregation of pubescent to glabrous plants, indicating the action of a single, dominant locus (we did not attempt to distinguish degrees of pubescence in this evaluation, and we do not exclude partial dominance or an additive effect). We selected two strongly pubescent and three strongly glabrous individuals for genotyping using the DArT-Seq platform. The pubescent individuals carried a large block of Mi21 introgression across chr 3 that was absent from glabrous plants (Fig. 3-S1). Introgression carried in the pubescent NIL spanned the qMHP3 interval identified in the B73xPT population, providing an independent line of evidence for a QTL in this location. There was no evidence of significant Mi21 introgression on chr 7, 8 or 9 in these BC_5_S_1_ individuals, supporting our previous conclusion that macrohair QTL are individually sufficient to promote a degree of stem macrohair production. The BC_5_S_1_ family provides a good starting point towards fine mapping and cloning of qMHP3.

### Comparison of B73xPT QTL and broader landrace diversity

To compare QTL detected in our B73xPT BC_1_S_5_ population to the broader diversity present in Mexican highland maize, we performed an environmental-genome wide association study (eGWAS) using a previously genotyped panel of 1830 geo-referenced Mexican landrace accessions (Romero Navarro *et al.*, 2017; Gates *et al.*, 2019; Fig. 6A; 4-S1). We selected the top 1000 SNPs most significantly associated with elevation and compared their physical location with the location of our QTL. The strongest environmental association was detected on chromosome 4 at the previously reported inversion polymorphism *Inv4m* (Romero Navarro *et al.*, 2017; Crow *et al.*, 2020). Although qED4, detected in our QTL mapping, co-localized with this region on chromosome 4 (Fig. 6B), we found no signal linking *Inv4m* to additional yield components or evidence of a previously reported flowering time effect in our QTL analysis. Further high confidence SNPs were found on chromosomes 2, 3, 5, 7, 8 and 10 (Fig. S4). A high confidence SNP on chr 7 (7_17794242) was identified adjacent to the GEI peak associated with qEW7 (Fig. 6C). This region was also identified in a previous experiment to map yield and harvest index QTL in a comparison of lowland and highland Mexican maize (Jiang *et al.*, 1999). In other cases, for example qTKW8 (Fig. 6D), there was no strong correspondence between the eGWAS hits and the location of the QTL. Although we would not draw strong conclusions from differences between eGWAS and QTL results, those cases where they do overlap provide compelling candidates for functional study. For example, the SNP 7_17794242 lies within a gene (Zm00001d019117) encoding a putative transmembrane protein that has been shown in temperate maize to be differentially expressed in response to salt, cold and UV (Makarevitch *et al.*, 2015).

**Figure 6.**
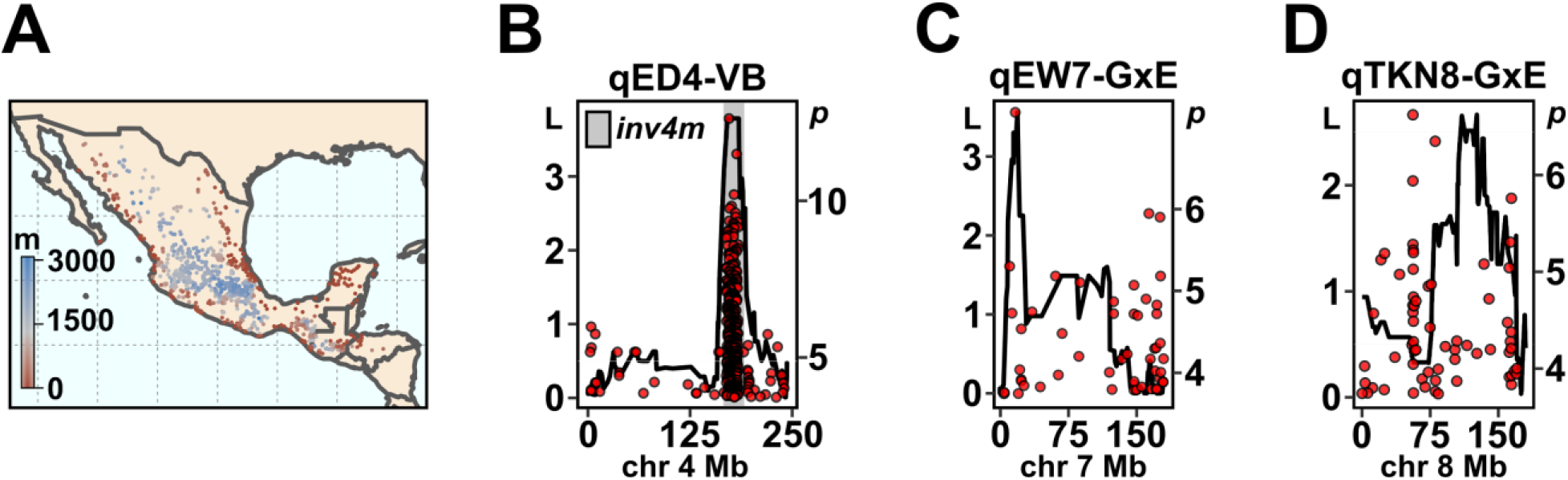
Colocalization of QTL with SNPs showing elevational variation in Mexican landrace maize. A) Geographic distribution of Mexican maize landraces. The color gradient represents elevation of the associated sampling location of maize landrace accessions. B) Trace showing support (LOD, L) for an *ear diameter* (ED) QTL across chromosome 4 (chr 4; physical distance) and SNPs (red points) significantly (-log_10_*p*, *p*) associated with elevation in Mexican maize landraces. LOD profile drawn using physically anchored genetic markers and trait values from the lowland site (VB). The gray rectangle indicates the position of the previously characterized *inv4m* inversion polymorphism. C, D) as B, showing support for *ear weight* (EW) and *total kernel number* (TKN) QTL on chromosomes 7 and 8, respectively. LOD profiles associated with trait GEI (GxE) values.

## DISCUSSION

Evaluation of a B73xPT mapping population in lowland and high elevation field sites identified QTL associated with both morphological and yield components traits. Although showing plasticity, the genetic architecture of morphological traits was conserved across environments and we saw little evidence of GEI. Indeed, characteristic highland traits such as pigmentation and pubescence were actually easier to evaluate in the lowland field as a result of the overall greater vigor of the plants. In contrast, we saw greater evidence of GEI with respect to yield components, with individual BC_1_S_5_ genotypes showing the signature of local adaptation and others stability across our two test environments. This broad trend of greater stability of morphological traits rather than yield components is consistent with a previous study mapping maize adaptation across four elevations (Jiang *et al.*, 1999).

In total, across the different phenotypic sets, we detected 44 unique QTLs, eighteen of which present a significant QEI where the majority (17) are examples of conditional neutrality, while only one (qEW7) was associated with statistical support for antagonistic pleiotropy. In a review of genetic architecture in 37 studies, the authors estimated that ∼60% of the QTLs detected displayed QEI, but that there was only evidence for antagonistic pleiotropy in ∼2% of cases (Des Marais *et al.*, 2013). This broad trend was reflected in a recent multisite mapping experiment in switchgrass (*Panicum virgatum L.*) in which the majority of QTL associated with adaptive traits showed conditional positive effects in their home environment with little or no detectable effect or cost in other environments (Lowry *et al.*, 2019). Detecting antagonistic pleiotropy requires higher statistical power than identification of conditional effects (*i.e.* the latter are typically supported by a failure to detect an effect in certain environments), resulting in a potential bias in the classification of QEI (Anderson *et al.*, 2011). In our study, the statistical power necessary to dissect QEI is limited by the size of the mapping population and the number of trials and locations evaluated. QTL effects for biomass, yield components changed in both magnitude and direction over location, suggesting antagonistic pleiotropy, and additional evaluation of our material may provide more evidence of this. We would also point out the extensive nature of the management employed at the lowland (Valle de Banderas) site; this site might be considered as an “ideal” environment in comparison to the highland site, exposing the plants to few of the stresses traditionally faced in tropical lowland fields. Any buffering of the potential costs of highland variants in the lowland site by management would push the genetic architecture from antagonistic pleiotropy towards conditionality.

We identified several QTLs that could be confidently associated with strong candidate genes. For stem pigmentation, qPINT2 and qPINT10 correspond well to the loci *b1* and *r1*, respectively. The qPINT2 locus had the greatest effect of the two (explaining ∼40% variance) with the PT allele promoting pigmentation. The *b1* gene encodes a bHLH transcription factor that regulates the temporal and tissue-specific expression of genes that produce anthocyanins in maize (Ludwig *et al.*, 1989; Petroni *et al.*, 2000; Chatham & Juvik, 2021). Interestingly, *b1* was also identified in a mapping cross between lowland and highland teosinte, the latter showing the stem pigmentation also seen in highland maize (Lauter *et al.*, 2004). Several independently derived *B1* alleles have been linked to stem pigmentation (Dooner & Kermicle, 1976; Radicella *et al.*, 1992; Selinger *et al.*, 1998; Selinger & Chandler, 1999, 2001; Chatham & Juvik, 2021), indicating the ready production of functional diversity at this locus and implicating convergent selection (Stern, 2013) for pigmentation among highland *Zea*, supporting an adaptive role (Doebley, 1984; Lauter *et al.*, 2004). Further sequencing of *b1* alleles from highland maize will shed greater light on patterns of diversity and the origin of different alleles. Stem pigmentation, unlike stem pubescence, is also shared with South American highland maize (Janzen *et al.*, 2021). Dark red pigmentation in the stem can help the plant to absorb more solar radiation and keep the plant warmer in a cold environment and might also protect DNA from damage due to higher UV-B radiation in the highlands (Barthakur, 1974; Eagles & Lothrop, 1994; Casati & Walbot, 2005) - although it is unclear why such protection might be required more so in the stem than in the leaf blades.

The flowering QTL qDTA8, qDTS8 and qASI8 overlap a ∼10Mb region that contains the two well-characterized flowering genes *Vgt1* and *Zcn8*. This region and/or these genes have been reproducibly detected in linkage- and association-mapping studies of maize flowering time (Chardon *et al.*, 2004; Buckler *et al.*, 2009; Steinhoff *et al.*, 2012; Li *et al.*, 2016; Romero Navarro *et al.*, 2017), temperate adaptation (Ducrocq *et al.*, 2008; Bouchet *et al.*, 2013; Guo *et al.*, 2018; Castelletti *et al.*, 2020) and adaptation to the Mexican Highlands (Gates *et al.*, 2019; Janzen *et al.*, 2021; Wang *et al.*, 2021). An early flowering *vgt1* allele from northern germplasm has previously been associated with a miniature transposon (MITE) insertion, although the absence of the MITE alone did not explain late flowering *vgt1* alleles (Buckler *et al.*, 2009). In a *Zcn8* association study using maize and teosinte, the haplotype associated with earliest flowering (A-Del) was hypothesised to have originated in highland teosinte (*Zea mays* ssp. *mexicana*) and to have moved to cultivated maize by introgression (Guo *et al.*, 2018). Interestingly, in this same study the authors report Palomero Toluqueño to carry both the MITE-associated allele of *vgt1* and the A-Del haplotype of *Zcn8*. Although we have not sequenced the *Vgt1* and *Zcn8* alleles present in our mapping population and available genome sequence data do not provide good coverage in this region, our linkage mapping results are consistent with this previous association analysis.

We identified QTLs associated with sheath pubescence on chr 3, 7, 8 and 9. Our QTL on chr 9 is consistent with previous observations co-localizing i) a leaf blade macrohair mutation in temperate maize, ii) a stem pubescence QTL in *mexicana* teosinte, iii) introgression from *mexicana* to highland maize and iv) a ∼ 3 Mb inversion that displays patterns of selection in mexican highland maize (Moose *et al.*, 2004; Lauter *et al.*, 2004; Hufford *et al.*, 2013; Gonzalez-Segovia *et al.*, 2019; Calfee *et al.*, 2021). Identification of a PT allele linked to stem pubescence in this same region of chr 9 adds further support to the hypothesis of adaptive introgression at this locus (Wilkes, 1972; Gonzalez-Segovia *et al.*, 2019). In this context, it is interesting to note that all macrohair QTL identified appeared to be sufficient on their own to induce stem pubescence, although their combined action would be needed to approach the level of pubescence of the PT parent. Limitations of population size and the semi-quantitative nature of our evaluation prevent strong conclusions concerning additivity or interactions among macrohair QTL. Nonetheless, our data suggest that qMH9 is just one of a number of contributing loci, the origin of the trait being a complex mix of wild-relative introgression and *de novo* mutation. Fine mapping and molecular cloning of the genes underlying macrohair QTL would allow a far more detailed view of the history of the stem pubescent trait and associated genetic variants in Mexican highland maize. Pubescence extends the boundary layer around the stem and could act as protection from cold by preventing heat loss or conserve water by minimizing transpiration (Chalker-Scott, 1999; Schuepp, 1993). We did not observe any strong correlation between pubescence and yield components. However, further experiments making use of the range of pubescence in our inbred families or derived NILs, might have the power to detect more subtle effects in either controlled conditions or large-scale highland yield trials.

The rich diversity of Mexican landrace maize is closely tied to local adaptation. Yet, this same specialization places these varieties at risk from future climate change (Mercer & Perales, 2010)(Bellon *et al.*, 2011) (Romero *et al.*, 2020). In a study to project landrace distribution under different climate change scenarios, PT was the landrace identified as the most vulnerable (Ureta *et al.*, 2012), although, as the authors note, models based on current distribution and climate do not take into account the full range of environmental, biotic and cultural factors that impact diversity and distribution. In the specific case of PT, limited yield potential in comparison to more modern landraces is likely to see it abandoned by farmers (https://www.biodiversidad.gob.mx/diversidad/proyectoMaices). That said, PT has contributed to the broader Mexican highland group (Reif *et al.*, 2006; Warburton *et al.*, 2008)(Arteaga *et al.*, 2016) and locally-adapted alleles will likely be conserved. The fate of the Mexican highland maize group will be influenced by their degree of resilience and ability to adapt to climate change. Although we found some evidence of antagonistic pleiotropy, PT and PT allele effects were largely stable and GEI was driven by plasticity associated with B73. As such, our data would support cautious optimism that highland varieties might maintain current levels of productivity in the face of future climate change. However, if climate change results in the expansion of lower elevation varieties to the highlands, the home site advantage of traditional highland landraces might be eroded. Ultimately, the conservation of maize diversity, along with the responsible and equitable use of this unique resource, will be informed by a greater understanding of the physiological and mechanistic basis of local adaptation. With an increase in mapping resolution (for example by using larger or more diverse populations) and the availability of high quality landrace genome assemblies, it will be possible to take important further steps towards defining not just the genetic architecture but also the genes and genetic variants that underlie local adaptation in maize landraces.

## FUNDING SOURCES

Consejo Nacional de Ciencia y Tecnología [Mexico] (FOINS-2016-01)

Consejo Nacional de Ciencia y Tecnología [Mexico] (CB-2015-01 254012)

Secretaria de Agricultura y Desarrollo Rural (Ministry of Agriculture and Rural Development: SADER) of the Government of Mexico under the MasAgro (Sustainable Modernization of Traditional Agriculture) initiative

National Science Foundation [USA] (No. 1546719)

UC-MEXUS (CN-15-1476)

RJHS is supported by the USDA National Institute of Food and Agriculture and Hatch Appropriations under Project #PEN04734 and Accession #1021929.

---

**Table S1:** Environmental characteristics of the experimental sites in Metepec (MT) and Valle de Banderas (VB) located in Mexico.

| **Field site** | **Valle de Banderas (VB)**<br>**Lowlands** | **Metepec (MT)**<br>**Highlands** |
| --- | --- | --- |
| **Elevation (m.a.s.l.)** | **54** | **2610** |
| **State** | **Nayarit** | **Mexico State** |
| **Latitude** | **20.784** | **19.223** |
| **Longitude** | **-105.244** | **-99.547** |
| **Temperature**<br>**Min/Mean/Max (°C)** | **22.5/25.8/28.4** | **12.3/12.4/17.1** |
| **Precipitation (mm)** | **1173** | **809** |
| **Type of soil** | **Regosol** | **Andosol** |

**Figure S1 - Supplemental 1.**
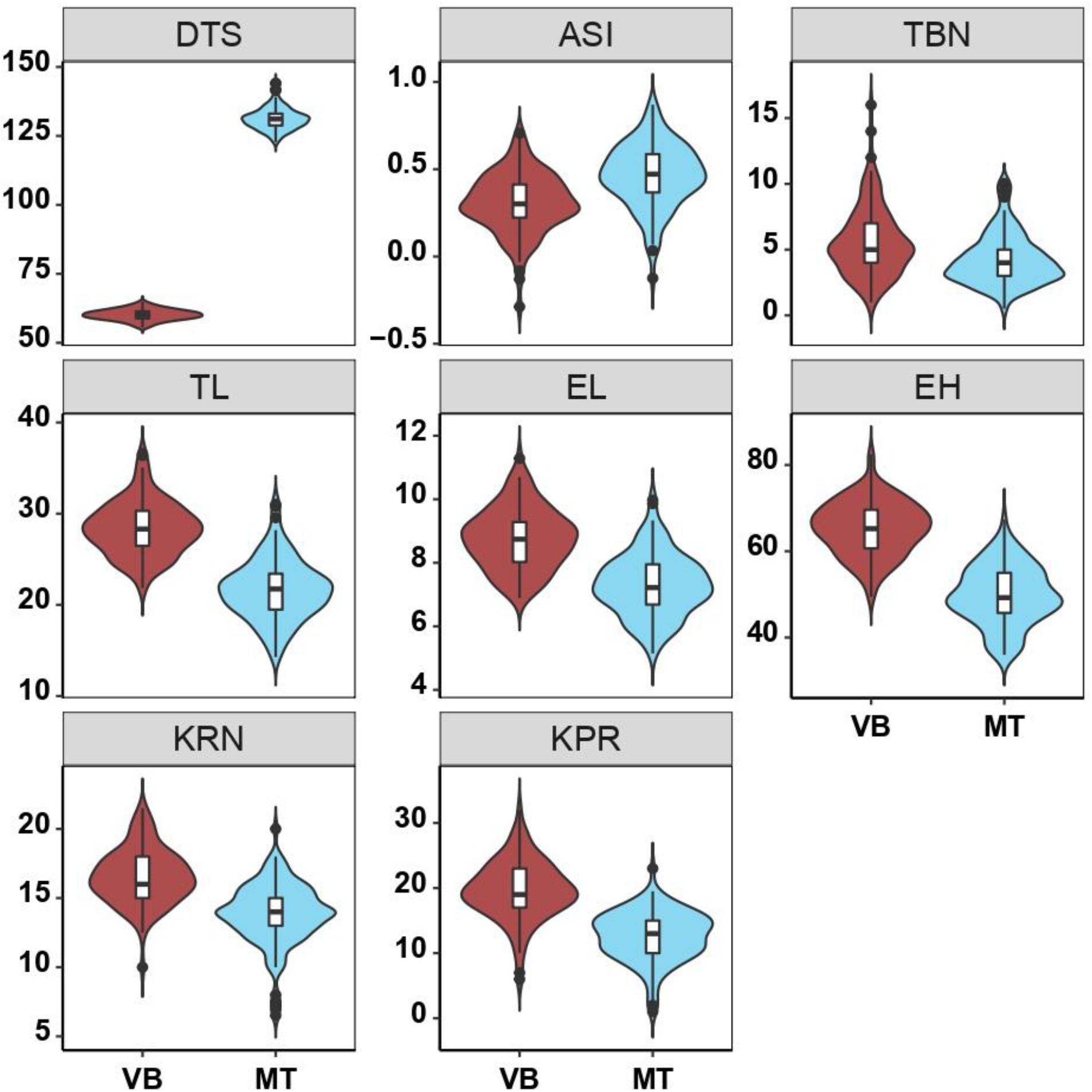
Distribution of plant phenotypic traits for B73xPT recombinant inbred lines grown in VB or MT field sites (trait descriptions were shown in Table 1). Fitted values for each genotype and location combination were used to generate violin plots and were estimated by adding BLUPs of the G+GEI to the estimated location mean. Median values were used to generate the violin plots for TBN, KPR, KRN. Boxes represent the interquartile range with the horizontal line representing the median and whiskers representing 1.5 times the interquartile ranges. The shape of the violin plot represents probability density of data at different values along the y-axis.

**Figure S2 - Supplemental 1.**
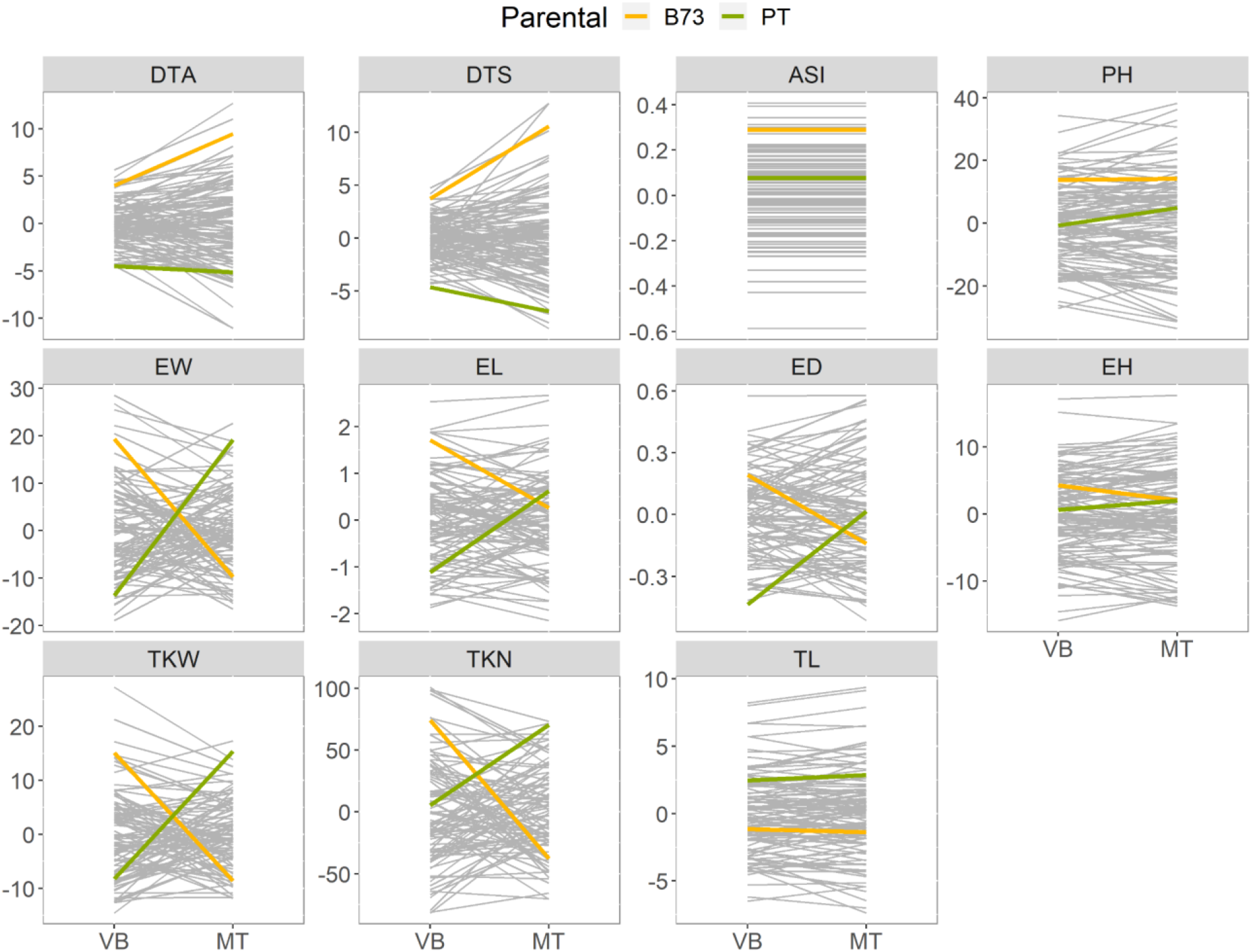
Reaction norm plots of plant phenotypic traits for B73xPT recombinant inbred lines grown in VB or MT field sites (trait descriptions were shown in Table 1). Values shown are the sum of Best Linear Unbiased Prediction (BLUP) for genotype and genotype by location interaction effects for each genotype in two field sites. Gray line segments connect values for each RIL genotype in the two field sites. B73 and PT parental values are shown in thick yellow and green lines respectively.

**Figure S3.**
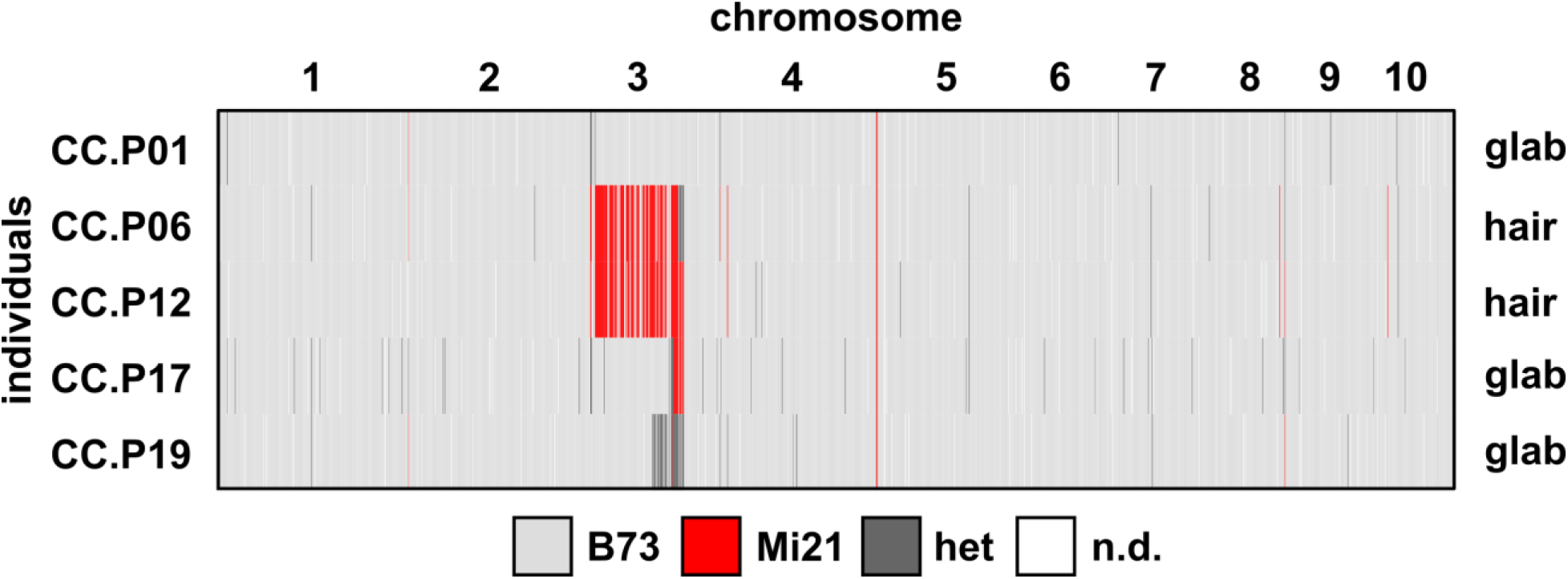
Pubescent segregants from a B73xMi21 BC_5_S_1_ family contain Mi21 introgression on chromosome 3. Three glabrous (glab) and two pubescent (hair) individuals genotyped with DaRT-SEQ.

**Figure S4.**
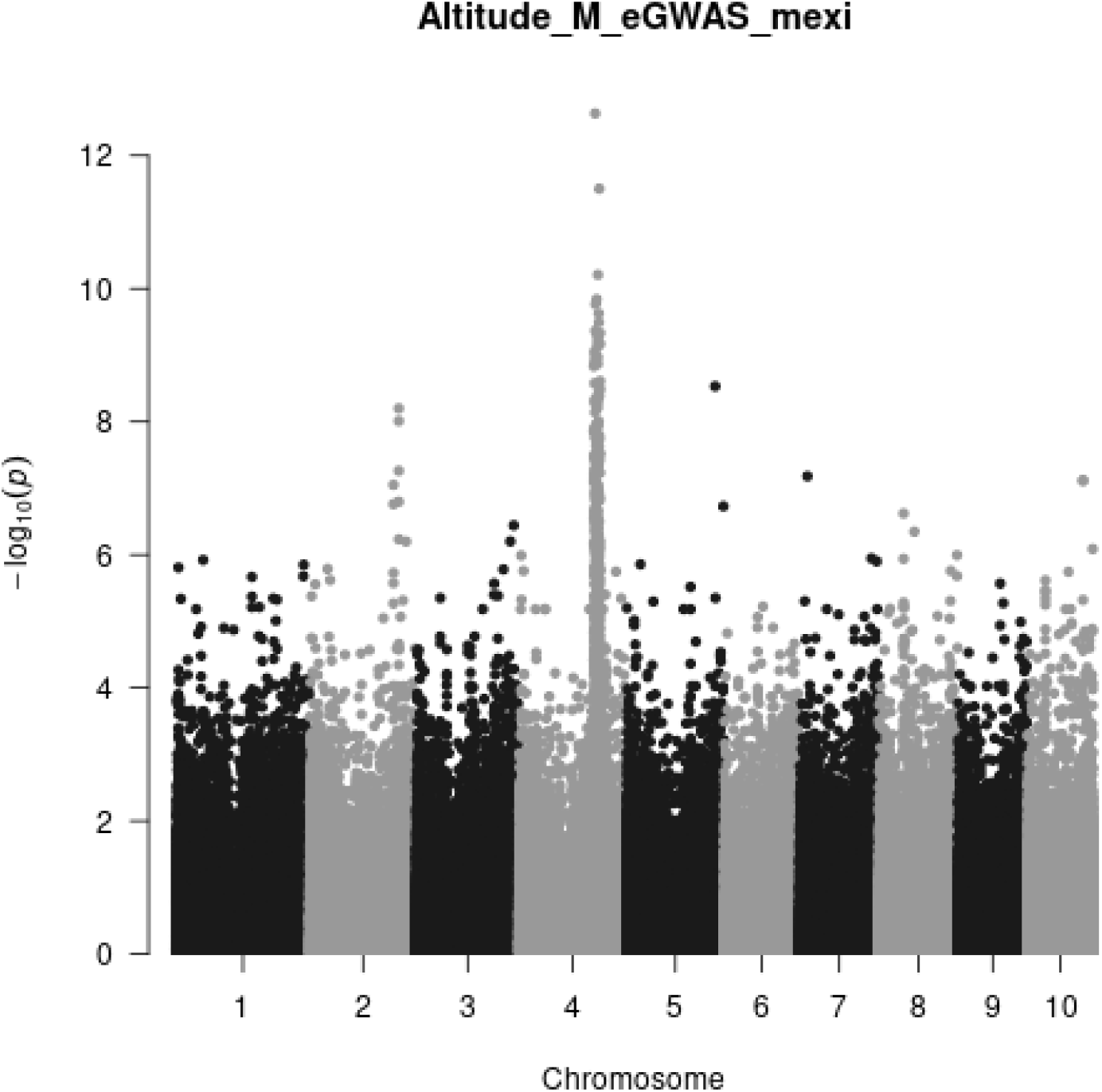
Elevation eGWAS Manhattan plot.

**Figure S5 - Supplemental 1:**
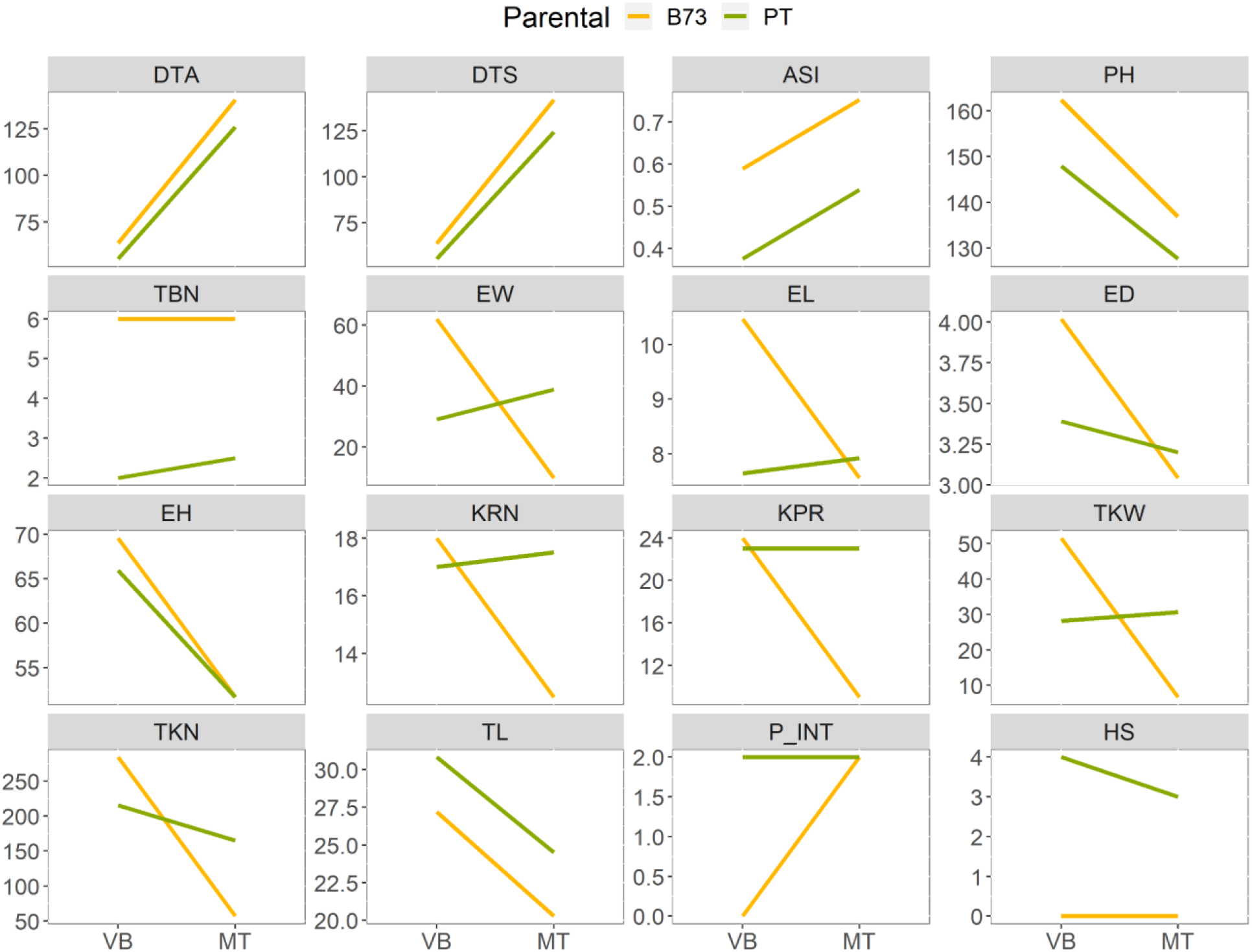
Reaction norms of B73 (green line) and Palomero Toluqueño landrace (yellow line) grown in VB and MT field sites (trait descriptions were shown in Table 1). The fitted values were estimated by adding BLUPs of the G+GEI to the estimated location mean. Palomero Toluqueño values were obtained by evaluating multiple heterozygous individuals from MEXI5 accession from CIMMYT.

**Figure 8 - Supplemental 1:**
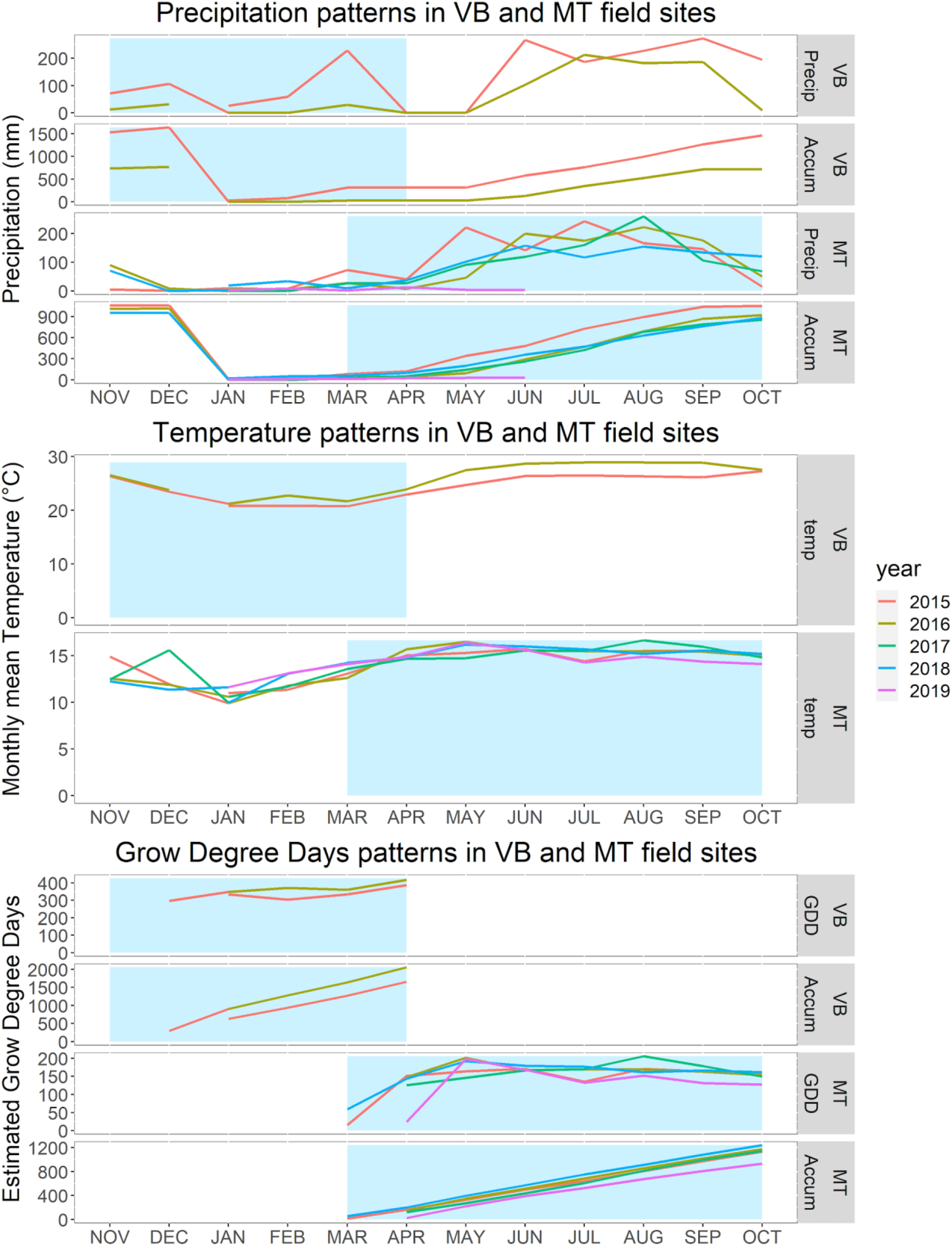
Monthly and accumulated precipitation, monthly average temperature and monthly and accumulated estimated Grow Degree Day patterns for VB and MT field sites for the years where evaluation was performed (2014-2015, 2015-2016 for VB field site; 2015, 2016, 2018 and 2019 for MT field site). The precipitation (in mm) and monthly mean temperature (°C) data was obtained from CONAGUA (National Commission on Water) database (https://smn.conagua.gob.mx/es/climatologia/informacion-climatologica/informacion-estadistica-climatologica; estation #15266 for MT site; estation #18030 for VB site) and temperature data was complemented with https://wu-next-ibm.wunderground.com/ where needed. Grow Degree Days estimation was performed with the formula GDD = (Tmean - 10)*n where Tmean is the monthly average temperature and n is the number of days accumulated in that month. The blue areas describe the typical growing season in winter in VB site (november-april) and summer in MT site (march-october).

